# Spindle pole body-associated Atg11, an autophagy-related protein, regulates microtubule dynamics essential for high-fidelity chromosome segregation

**DOI:** 10.1101/2021.12.15.472744

**Authors:** Md. Hashim Reza, Rashi Aggarwal, Jigyasa Verma, Nitesh Kumar Podh, Ratul Chowdhury, Gunjan Mehta, Ravi Manjithaya, Kaustuv Sanyal

## Abstract

Emerging studies hint at the roles of autophagy-related proteins in various cellular processes in a eukaryotic cell. To understand if autophagy-related proteins influence genome stability, we examined a cohort of 35 autophagy mutants in *Saccharomyces cerevisiae.* We observed cells lacking Atg11 show poor mitotic stability of minichromosomes. Atg11 molecules dynamically localize to the spindle pole bodies (SPBs). Loss of Atg11 leads to a delayed cell cycle progression. Such cells accumulate at metaphase at an elevated temperature that is relieved when the spindle assembly checkpoint is inactivated. Indeed, *atg11*Δ cells have stabilized securin levels, that prevent anaphase onset, confirming chromosome biorientation defects associated with the mutant. Atg11 functions in the Kar9-dependent spindle positioning pathway and maintains Kar9 asymmetry by facilitating proper dynamic instability of astral microtubules (aMTs). Taken together, this study uncovers a non-canonical role of Atg11 in facilitating MT dynamics crucial for chromosome segregation.

## Introduction

Dynamic interactions between spindle microtubules (MTs) and chromosomes facilitate precise segregation of the genetic material during the cell cycle. Comprising of α/β-tubulin dimers, MTs display inherent polarity, with the minus end proximal to the spindle poles, while the more dynamic plus end interacts with the chromosomes at the kinetochore. *Saccharomyces cerevisiae* and related budding yeast species undergo closed mitosis with an intact nuclear envelope throughout the cell cycle (*1–5*). Spindle pole bodies (SPBs), the microtubule organizing centers (MTOCs), in these organisms, remain embedded in the nuclear envelope throughout the cell cycle (*6*) and emanate both nuclear and astral microtubules (aMTs). Some of the nuclear MTs attach to kinetochores and facilitate the poleward movement of sister chromatids required for the precise segregation of chromosomes in progeny daughter cells. The spindle assembly checkpoint (SAC), which detects any defects in kinetochore-MT attachments or improper tension at kinetochores, halts the cell cycle at metaphase until all errors are corrected and prevents any premature separation of sister chromatids (*7, 8*). The SAC inhibits the anaphase-promoting complex/cyclosome (APC/C), a ubiquitin ligase required for the degradation of securin and subsequent release of separase, that is necessary for cohesin cleavage and sister chromatid separation (*9*).

The cytosolic face of SPBs nucleates aMTs which aid in the alignment and positioning of the mitotic spindle along the polarity axis, critical for proper nuclear division during asymmetric cell division in *S. cerevisiae*. Two pathways ensure asymmetric nucleation of aMTs from the old SPB (dSPB) in yeast. The cyclin-dependent kinase (CDK) Cdc28/Cdk1, representing the intrinsic factor, facilitates the biased recruitment of SPB components such as Spc72 and the γ-tubulin complex (γ-TC) at the dSPB, while asymmetric localization of Kar9 at the dSPB, represents the extrinsic factor (*10–12*). Such asymmetry leads to the nucleation of more aMTs from the dSPB necessary for the mitotic spindle positioning along the mother-bud axis and ensures non-random SPB inheritance (*12–18*). Misaligned spindles activate the spindle position checkpoint (SPOC) (*19–23*), a surveillance mechanism that halts the cell cycle and prevents exit from mitosis by inhibiting the mitotic exit network (MEN) signaling pathway (*24*).

The stochastic transitions between growth and shrinkage, called dynamic instability, are exhibited by MTs in the cell (*25*). MT dynamic instability is defined by four parameters: frequency of catastrophe (the transition from polymerization to depolymerization), frequency of rescue (the transition from depolymerization to polymerization), and the rates of polymerization and depolymerization (*26*). The dynamic instability facilitates MTs to search for the binding sites on kinetochores (the search-capture model) and thereby helps in the initial attachment of chromosomes to the mitotic spindle (*27*). It also aids in the interaction of aMTs with the cell cortex, crucial for spindle positioning and alignment (*13*). Proteins regulating MT dynamics have been classified as either MT-stabilizing factors or MT-associated proteins (MAPs) which regulate MT polymerization or MT destabilizing factors which induce catastrophe (*26*). A proper balance of these regulatory factors ensures controlled MT dynamics facilitating proper and timely nuclear division in the cell. Therefore, any perturbations in MT dynamics have wider implications ranging from failure in embryonic development, early ageing, and aneuploidy, to cancer (*28–32*). Therefore, it is imperative to study components that preserve MT integrity crucial for both positioning of the nucleus and proper kinetochore-MT attachments during the cell cycle.

Autophagy, apart from being a conserved cellular degradation process, is also implicated in cell division under stress conditions (*33, 34*) and in controlling the integrity of nuclear-envelope-embedded nuclear pore complexes during the cell cycle (*35, 36*). A series of emerging evidence highlights the role of autophagy in cell metabolism, growth, aging, and genome stability (*33, 34, 37*). Under stress and starvation conditions, autophagy is induced and involves Atg17-meditated non-selective degradation of cargoes, while Atg11 mediates selective autophagy predominantly under vegetative conditions (*38*). In this study, we screened a collection of autophagy mutants and identified Atg11 as a regulator of high-fidelity chromosome segregation in *S. cerevisiae*. Our results uncover the previously unknown role of Atg11 in regulating MT integrity critical for proper kinetochore-MT interactions. We provide evidence that a significant cellular pool of Atg11 is SPB-associated ensuring proper cell cycle stage-specific dynamics of aMTs vital for nuclear positioning.

## Results

### Atg11 plays a role in high-fidelity chromosome transmission

To understand the role of autophagy protein(s) in the cell cycle in nutrient-rich conditions, we carried out a phenotypic screen to determine the growth fitness of null mutants of autophagy-related genes (*atg*) in *S. cerevisiae* to a microtubule depolymerizing drug, thiabendazole (TBZ). Among all the 35 *atg* mutants (f**ig. S1A**), *atg11*Δ cells displayed maximum sensitivity to TBZ (**Fig. 1A**), supporting an earlier observation (*39*).

**Fig S1.**
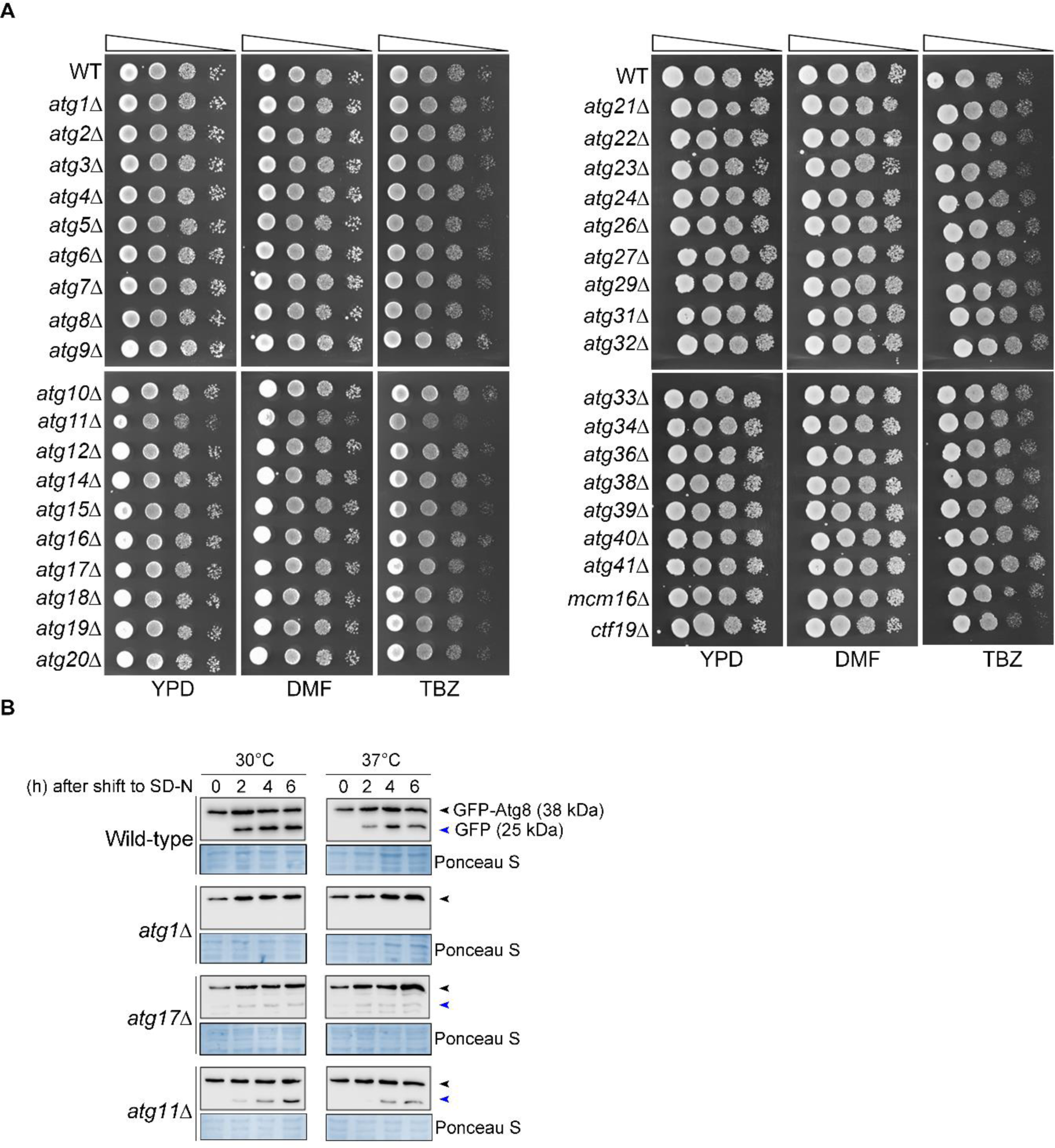
The *atg11*Δ cells are hyper-sensitive to the microtubule depolymerizing drug thiabendazole (TBZ). (A) Overnight grown cells of wild-type, null mutants of autophagy-related genes (*atg*), and two kinetochore mutants, *mcm16*Δ and *ctf19*Δ, were 10-fold serially diluted, spotted on YPD and YPD plates containing dimethylformamide (DMF) only or 50 μg mL^-1^ thiabendazole (TBZ). Plates were photographed after incubation at 30°C for 36 h. (B) Immunoblot analysis of whole-cell lysate prepared at indicated time points from the wild-type, *atg1*Δ, *atg17*Δ, and *atg11*Δ strains expressing GFP-Atg8 grown at 30°C and 37°C in nitrogen starvation medium and probed with anti-GFP antibodies. GFP-Atg8 (38 kDa) and free GFP (25 kDa) are labeled with black and blue arrowheads, respectively.

**Fig 1.**
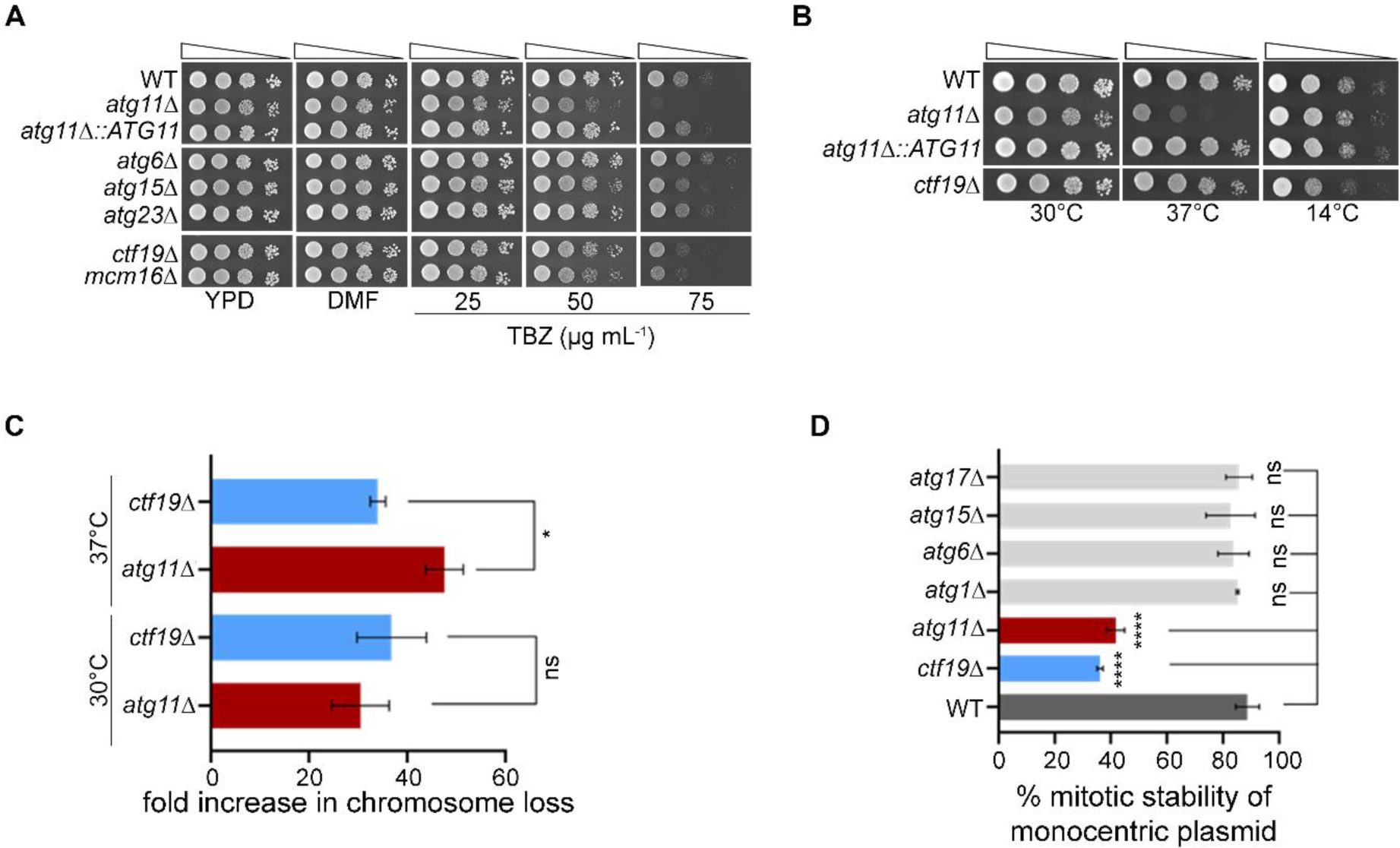
Cells lacking Atg11 show an increased rate of chromosome loss. (A) Overnight grown cells of wild-type, null mutants of autophagy-related genes (*atg*), and two kinetochore mutants, *mcm16*Δ and *ctf19*Δ, were 10-fold serially diluted, spotted on YPD and YPD plates containing dimethylformamide (DMF) only or 25, 50, 75 μg mL^-1^ thiabendazole (TBZ). Plates were photographed after incubation at 30°C for 48 h. (B) Overnight grown cells of indicated strains were 10-fold serially diluted and spotted on YPD plates. Plates were photographed after incubation at 30°C and 37°C for 48 h, or 14°C for 7 days. (C) Bar graph showing fold increase in chromosome loss in *atg11*Δ and *ctf19*Δ strains at 30°C and 37°C from three biological replicates. Error bars indicate standard deviation (SD). Statistical analysis was done using Ordinary one-way ANOVA using Tukey’s multiple comparisons tests (*p*=*0.0439). (D) Mitotic stability of a monocentric plasmid (pRS313) was determined in wild-type, *atg* mutants as indicated, as well as in the kinetochore mutant strain *ctf19*Δ. The assay was done with three independent sets of transformants. Statistical analysis was done using one-way ANOVA using Dunnett’s multiple comparisons tests (****p*<*0.0001).

Transgenic wild-type *ATG11* complemented the TBZ-induced growth defects in *atg11*Δ cells (**Fig. 1A**). Furthermore, *atg11*Δ cells grew as good as wild-type at 14°C but significantly slower at 37°C (**Fig. 1B**), a condition at which the rate of depolymerization of MTs is more dominant than the rate of polymerization (*40*). These results hint towards the role of Atg11 in regulating MT-dependent processes.

Any defects in MT-dependent processes negatively impact the fidelity of chromosome segregation. Indeed, a strain harboring an extra linear chromosome displayed a significantly higher rate of chromosome loss in *atg11*Δ cells at 30°C that was further exacerbated at 37°C (**Fig. 1C**). Similarly, the average mitotic stability of a monocentric circular plasmid was found to be significantly lower in *atg11*Δ cells than in the wild-type but comparable to that of a kinetochore mutant, *ctf19*Δ (**Fig. 1D**). No other *atg* mutants tested displayed any decrease in the mitotic plasmid stability. Mutants of key autophagy proteins such as Atg1, essential for both cytoplasm-to-vacuole targeting (Cvt) and autophagy pathways, or Atg17, required during starvation (*41*), neither displayed any TBZ sensitivity nor any reduction in plasmid stability (**fig. S1A** and **Fig. 1D**). Using GFP-Atg8 processing assay (*42*), as previously reported, the *atg1*Δ cells were defective for autophagy (absence of free GFP), while both *atg11*Δ and *atg17*Δ cells have reduced levels of autophagy relative to the wild-type (**fig. S1B**). This suggests that while Atg1 is indispensable for autophagy, it does not play any role in chromosome segregation as shown by our assays. Taken together, we conclude that the loss of Atg11 in yeast reduces the fidelity of chromosome segregation leading to an increased frequency of chromosome loss.

### Atg11 displays a dynamic association with the SPBs

To understand how Atg11 contributes to MT-dependent processes, we first studied the dynamics of Atg11 during the cell cycle progression. A protein-protein interaction network analysis of components involved in autophagy and chromosome segregation identified Spc72, one of the components of SPBs, as a key interactor of Atg11 (**fig. S2A**) (*43*), raising the possibility that Atg11 could be associated with the SPBs during the cell cycle.

**Fig S2.**
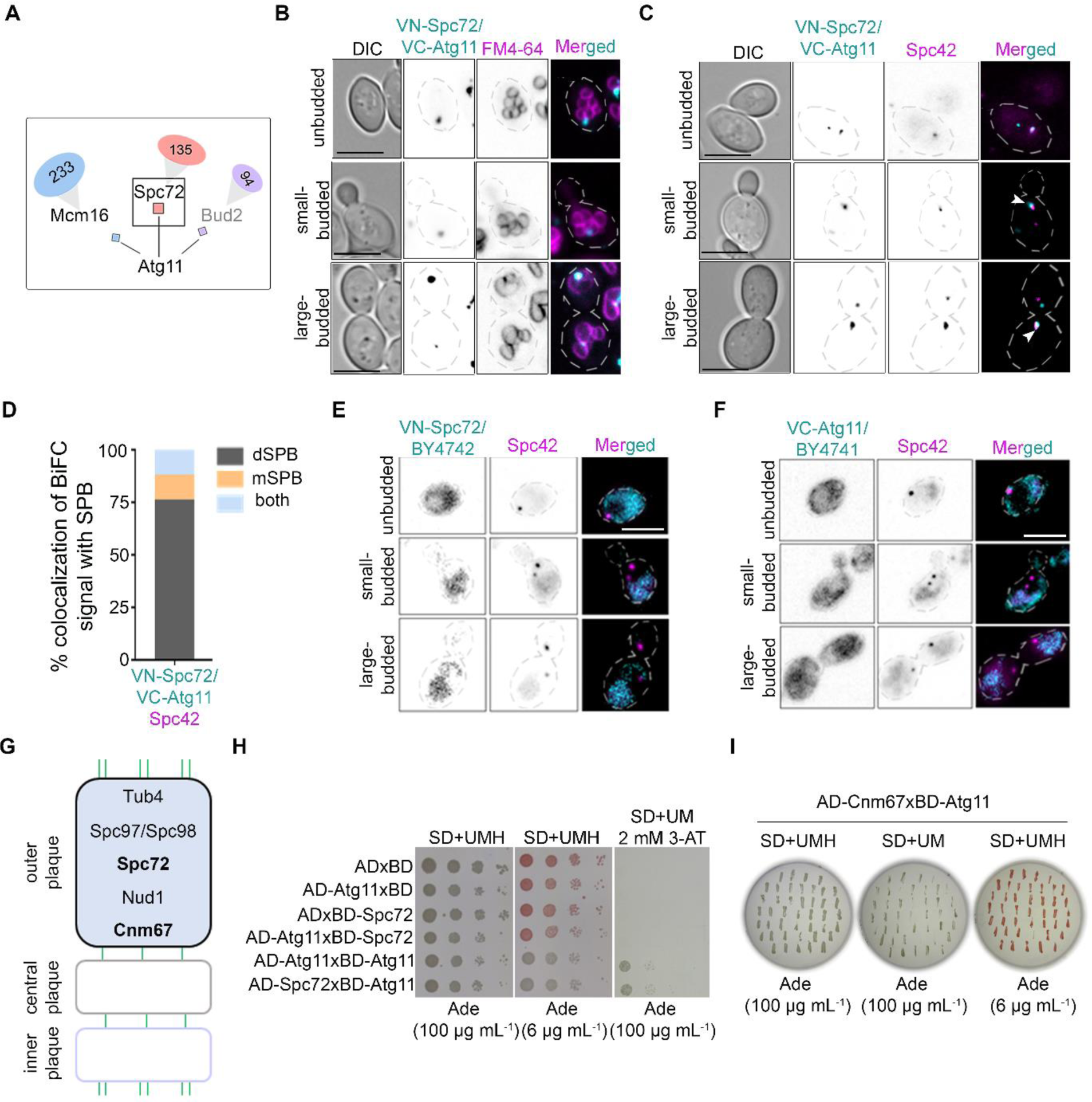
Atg11 interacts with Spc72 at the SPBs. (A) Primary (square boxes) and secondary (oval) interactors of Atg11 as reported in the latest curation of BioGRID (*43*). (B) Representative images displaying an *in vivo* BiFC interaction between Atg11 and Spc72 by the reconstitution of Venus fluorescence (VN-Spc72 /VC-Atg11) in diploid cells at the vacuolar periphery, marked by FM4-64 staining. Scale bar, 6 μm. (C) Representative images displaying an *in vivo* BiFC interaction between Atg11 and Spc72 at the SPBs, (Spc42-mCherry), in the diploid cells. The white arrowheads mark the co-localization of the BiFC signals with the dSPB. Scale bar, 6 μm. (D) A bar diagram displaying the proportion of cells displaying co-localization of BiFC signals with either dSPB, mSPB or both the SPBs (*n*=85 large-budded cells). (E) Representative fluorescence images expressing the VN-tagged Spc72 in diploid cells co-expressing Spc42-tagged with mCherry. Scale bar, 6 μm. (F) Representative fluorescence images expressing VC-tagged Atg11 in diploid cells co-expressing Spc42-tagged with mCherry. Scale bar, 6 μm. (G) Schematic showing the spatial position of proteins at the outer plaque of the SPB. (H) Yeast-two hybrid (Y2H) assays in *S. cerevisiae* strains carrying indicated plasmids were 10-fold serially diluted and spotted on SD - trp - leu or SD - trp - leu - his supplemented with 2 mM 3-amino-1,2,4-triazole (3-AT) and grown at 30°C. AD-activation domain, BD-DNA binding domain. (I) Yeast-two hybrid (Y2H) assays to study interactions between Atg11 and Cnm67, an outer plaque SPB protein (highlighted in bold in the schematic).

Live-cell imaging in a strain that co-expressed sfGFP-Atg11 and SPBs (Spc42-mCherry) revealed a transient and proximal localization of Atg11, first with the dSPB (14 out of 15 live-cell movies), before a cell enters anaphase (**Fig. 2A**).

**Fig 2.**
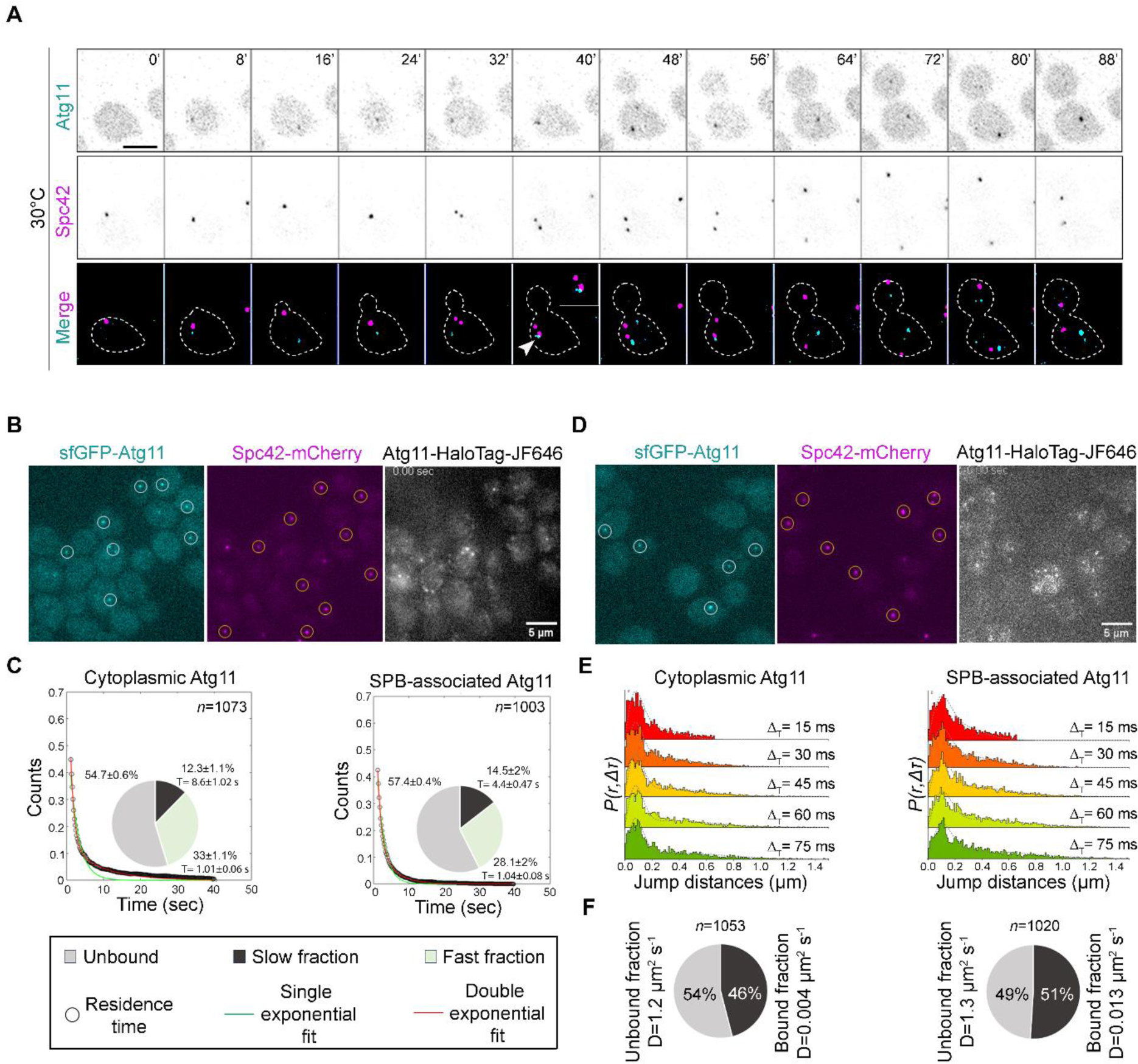
Single-molecule tracking quantifies the binding dynamics of Atg11 molecules at SPBs. (A) Time-lapse images showing Atg11 (sfGFP-Atg11) and SPB (Spc42-mCherry) dynamics during the cell cycle after G1 arrest followed by release at 30°C. The white arrowhead shows Atg11’s presence proximal to the dSPB. Scale bar, 5 μm. (B) Representative image for sfGFP-Atg11 puncta in the cytoplasm, Spc42-mCherry, and Atg11-HaloTag-JF646 imaged over 200 ms intervals. The regions of interest (ROIs) used for tracking Atg11 molecules present in the cytoplasm (cytoplasmic Atg11) or associated with SPBs (SPB-associated) are shown in white and orange circles, respectively. (C) Survival time distribution of Atg11 in the cytoplasm and SPB-associated was quantified from 200 ms time-interval movies. The distribution fits well with the double exponential curve, suggesting two types of bound population: 1) Fast fraction and 2) Slow fraction. Pie charts represent the percentage of molecules unbound (gray), bound with short residence time (fast fraction, light green), and bound with long residence time (slow fraction, black). The average residence times of fast and slow fractions are presented next to their representative fractions. *n*= number of tracks analyzed. (D) Representative image for sfGFP-Atg11 puncta in the cytoplasm, Spc42-mCherry, and Atg11-HaloTag-JF646 imaged over 15 ms intervals. The regions of interest (ROIs) used for tracking Atg11 over cytoplasmic Atg11 and SPBs are shown in white and orange circles respectively. (E) Spot-On based kinetic modelling for Atg11-HaloTag (obtained from 15 ms time interval movies). The probability density function histogram of single-molecule displacements for Atg11 in cytoplasm and over the SPBs is shown. The dashed line indicates the model derived from the Cumulative distribution function (CDF) fitting in Spot-On. (F) Pie charts represent the fraction of bound and unbound molecules with their mean diffusion coefficients (D), obtained from the Spot-On analysis. *n* = number of tracks analyzed.

To strengthen our observations of the transient localization of Atg11 to SPBs, we employed single-molecule tracking (SMT) to study the dynamic binding of Atg11 with SPBs. For the SMT analysis of Atg11, we endogenously tagged *ATG11* with a HaloTag at its C-terminus for the controlled labeling using HaloTag ligand Janelia Fluor 646 (JF646) (**Fig. 2B**) (*44*). We quantified the dwell time of Atg11 over SPBs (Spc42-mCherry) and Atg11 puncta (sfGFP-Atg11) in the cytoplasm. The time-lapse movies acquired with slow (200 ms time interval) and fast (15 ms time interval) imaging enabled the quantification of survival time and diffusion coefficient respectively. The cumulative survival time distribution fits well with the double exponential fit (**Fig. 2C**), suggesting two molecular populations: a) Atg11 puncta in the cytoplasm and b) Atg11 molecules associated with SPBs. We observed a significant two-fold decrease in the dwell time of Atg11 over Atg11 puncta (cytosol) versus SPBs (8.6±1.02 s vs. 4.4±0.47 s, respectively), suggesting a fast exchange of Atg11 over SPBs compared to the cytoplasmic puncta (**Fig. 2C**). We employed Spot-On-based kinetic modeling (*45*) to estimate the fraction of bound and unbound molecules with their mean diffusion coefficients (**Fig. 2D, E and F**). A marginal increase in the bound fraction of Atg11 over SPBs, compared to the cytoplasm (51% vs. 46%, respectively) (**Fig. 2F**) was observed. However, the diffusion coefficient of Atg11 over SPBs is three times higher than that of cytoplasm (D= 0.013 µm^2^/s vs. 0.004 µm^2^/s) (**Fig. 2F**), suggesting more dynamic interactions of Atg11 with SPBs, compared to the cytoplasmic puncta.

To support that Atg11 is transiently localized at the SPBs, we utilized a proximity-based bimolecular fluorescence complementation (BiFC) assay with Spc72, a known interactor of Atg11 (**fig. S2A**) (*46*). A fluorescent punctate signal resulting from the reconstitution of the Venus molecule at the vacuolar periphery, marked by FM4-64, suggested an *in vivo* interaction between Atg11 and Spc72 (**fig. S2B**). Co-localization with Spc42 further demonstrated the *in vivo* interaction between these two proteins at the SPBs (**fig. S2C and D**). The absence of BiFC signals in diploid cells expressing either VN-tagged Spc72 or the VC-tagged Atg11 alone (**fig. S2E and F**), suggested that neither VN nor VC fluoresces on its own. To further probe how Atg11 is associated with SPBs, we first confirmed a previously reported interaction between Atg11 and Spc72 (*47*) by the yeast two-hybrid (Y2H) assay (f**ig. S2G and H**). The Y2H assay with Cnm67, a protein from the outer plaque of the SPB, did not show any interactions with Atg11 (**fig. S2I**), suggesting the spatial position of Atg11 and Spc72 favors their physical interactions. These results, therefore, suggest a novel and dynamic association of Atg11 at the SPBs, in addition to its canonical localization onto the vacuolar membrane crucial for selective autophagy (*48*).

### Atg11 enables the timely progression of the cell cycle

SPBs nucleate both nuclear and astral microtubules, critical for kinetochore-MT interactions and spindle orientation along the mother-bud axis, respectively. Having established a non-canonical localization of Atg11 at the SPBs, we, therefore, looked at the status of both nuclear and aMT regulation in the absence of Atg11. First, we probed cell cycle progression in synchronized cells and studied the dynamics of tubulin (GFP-Tub1) and SPBs (Spc42-mCherry) in live cells. We observed a delay in spindle elongation and a concomitant delay in the cell cycle progression in *atg11*Δ cells (**Fig. 3A and B** and **fig. S3A and B**).

**Fig 3.**
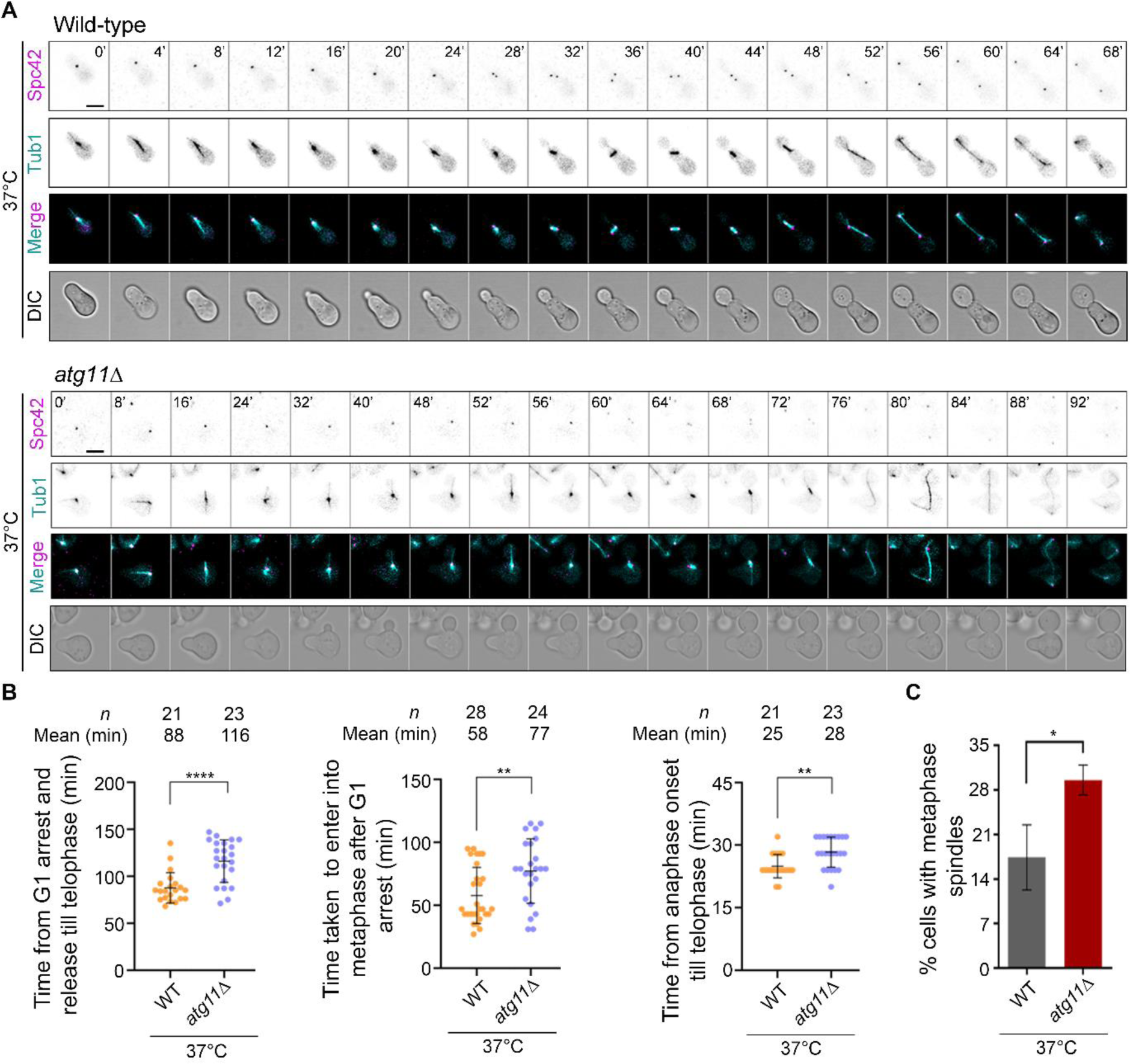
Cell cycle progression is significantly delayed in *atg11*Δ cells. (A) Time-lapse images showing dynamics of the mitotic spindle (GFP-Tub1) and SPBs (Spc42-mCherry) during the cell cycle in wild-type (*top*) and *atg11*Δ cells (*bottom*) after G1 arrest followed by release at 37°C. Scale bar, 5 μm. (B) Scatter plot displaying time taken for wild-type and *atg11*Δ cells, after G1 arrest followed by release at 37°C, for completion of the cell cycle (*left*), to enter into metaphase (*middle*, 1.5-2 µm mitotic spindle), and anaphase onset till telophase (*right*, disassembly of MTs). Error bars show mean ± SEM. Statistical analysis was done using an unpaired *t*-test with Welch’s correction (**p=0.0057/0.0011, ****p<0.001). (C) A bar graph representing the proportion of large-budded cells having metaphase (1.5-2 μm) spindle (GFP-Tub1) in wild-type and *atg11*Δ cells at 37°C. *n*>89, N=3. Error bars show mean ± SEM. Statistical analysis was done using an unpaired *t*-test with Welch’s correction (*p=0.0376).

**Fig S3.**
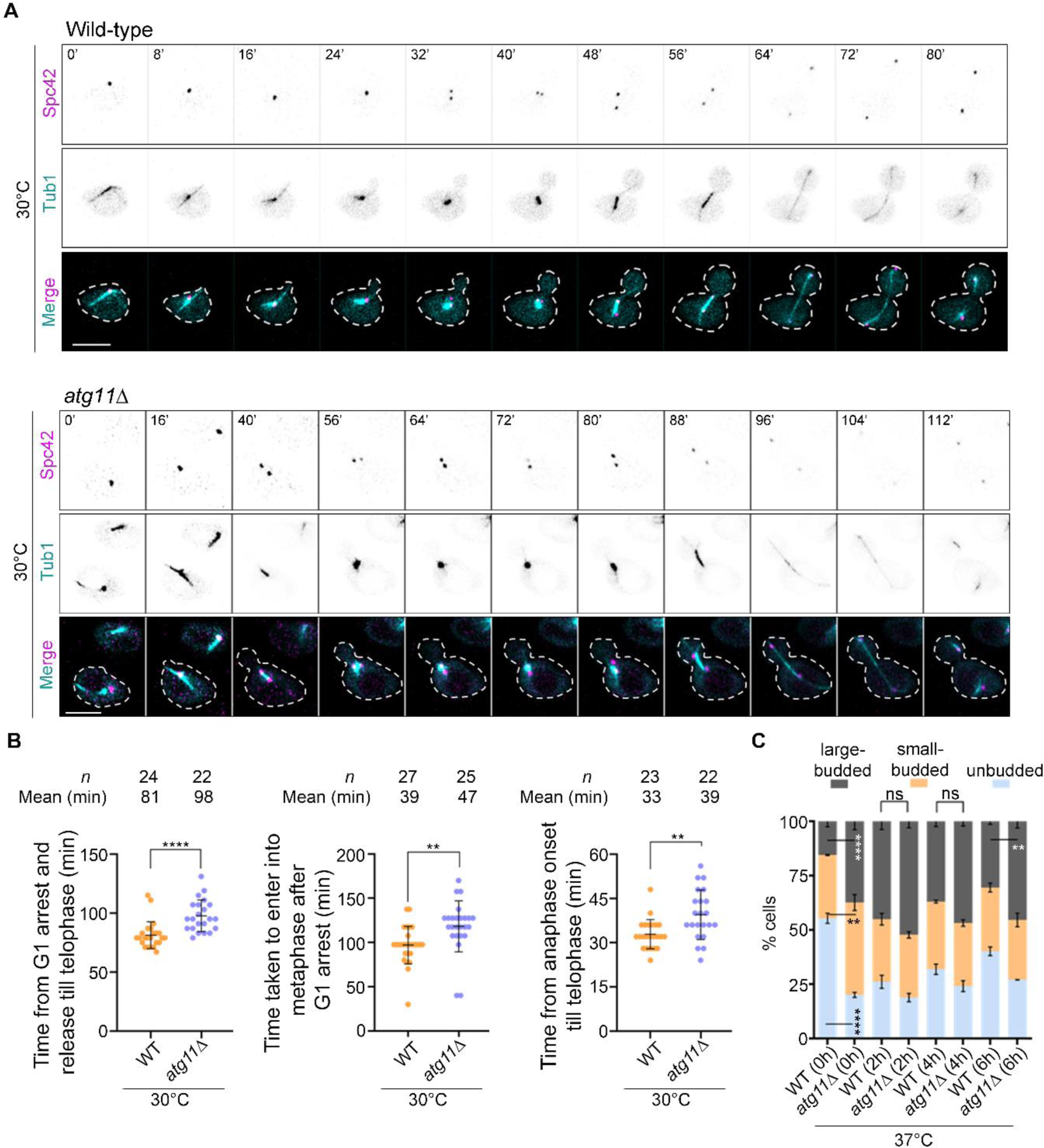
Cells lacking Atg11 display a delayed cell cycle at 30°C. (A) Time-lapse images showing dynamics of the mitotic spindle (GFP-Tub1) and SPBs (Spc42-mCherry) during the cell cycle in wild-type (*top*) and *atg11*Δ cells (*bottom*) after G1 arrest followed by release at 30°C. (B) Scatter plot displaying time taken for wild-type and *atg11*Δ cells, after G1 arrest followed by release at 30°C, for completion of the cell cycle (*left*), to enter into metaphase (*middle*, 1.5-2 µm mitotic spindle), and anaphase onset till telophase (*right*, disassembly of MTs). Error bars show mean ± SEM. Statistical analysis was done using an unpaired *t*-test with Welch’s correction (**p=0.0045/0.0031, ****p<0.001). (C) A bar diagram representing the proportion of unbudded, small-budded, and large-budded cells in wild-type (WT) and *atg11*Δ cells grown in YPD for 0, 2, 4, and 6 h at 37°C. More than 100 cells were analyzed for each biological replicate and for every time point, N=3. Error bars show mean ± SEM. Statistical analysis was done by two-way ANOVA using Tukey’s multiple comparisons test (**p=0.0068/0.0017, ****p<0.0001).

We also measured the time taken by *atg11*Δ cells to enter a) metaphase (spindle length=1.5-2 µm) after release from G1 arrest and b) telophase upon anaphase onset (spindle length>2 µm). We observed a significant delay at both these stages in *atg11*Δ cells (**Fig. 3B** and **fig. S3B**). Further analysis revealed that the overnight grown (t=0) *atg11*Δ cells displayed a significantly increased proportion of large-budded cells compared to the wild-type (f**ig. S3C**). When grown at 37°C for 6 h, the *atg11*Δ cells further accumulated an increased proportion of large-budded cells (**fig. S3C**), with a two-fold higher proportion of large-budded cells with metaphase spindle of 1.5-2 µm compared to the wild-type at 37°C (**Fig. 3C**), hinting towards a delay in anaphase onset.

Defects in kinetochore-MT attachments lead to the activation of the spindle assembly checkpoint (SAC), an error correction mechanism that delays the metaphase-to-anaphase transition (*49*). We, therefore, examined whether the SAC sensed the *atg11*-associated defects of delayed anaphase onset. Deletion of *MAD2* in the *atg11*Δ cells alleviated the proportion of large-budded cells with unsegregated nuclei demonstrating that indeed the SAC senses the *atg11*Δ-associated defects leading to delayed anaphase onset (**Fig. 4A**).

**Fig 4.**
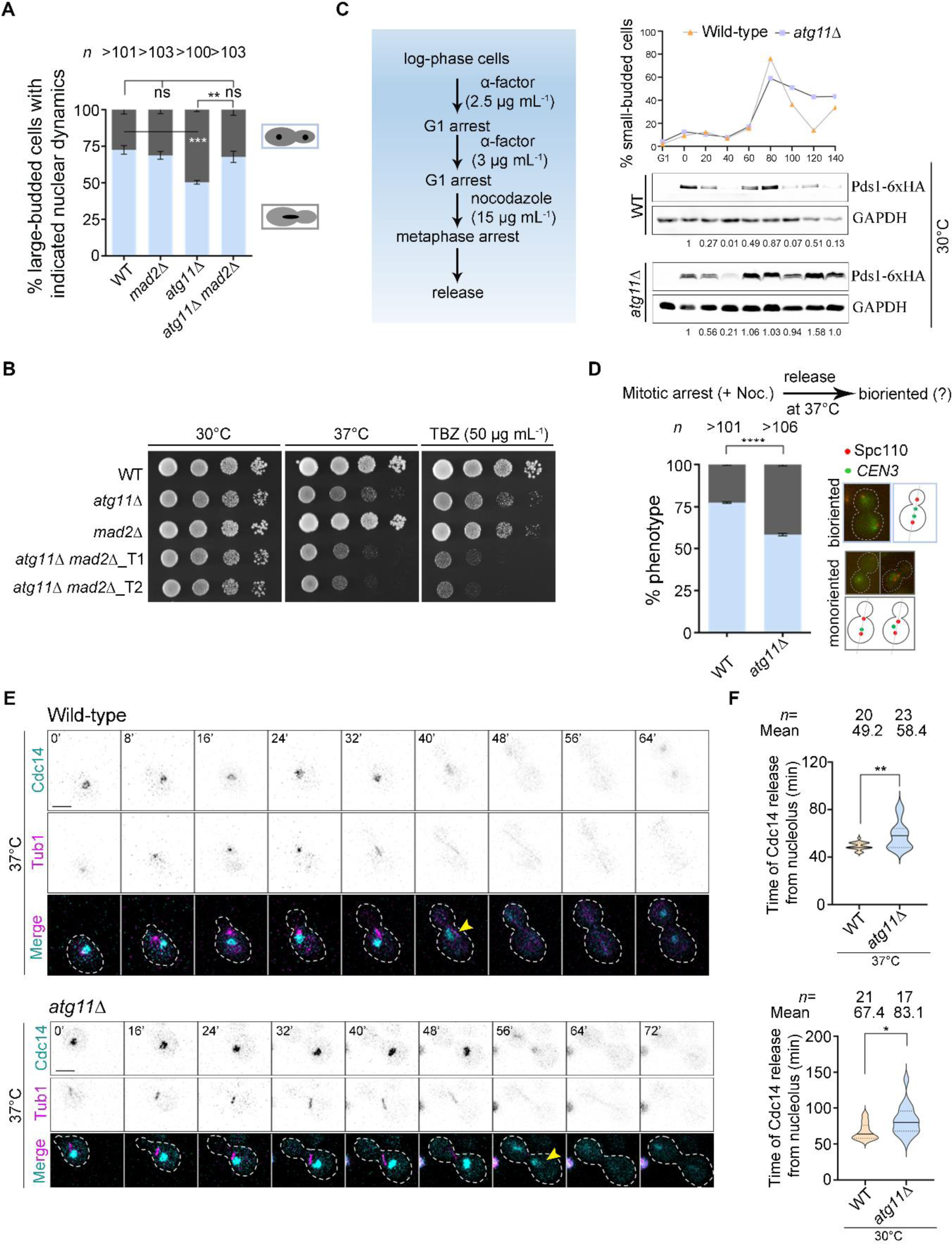
Mad2-mediated spindle assembly checkpoint delays cell cycle progression in *atg11*Δ cells. (A) A bar diagram representing the proportion of large-budded cells with stages of nuclear division in indicated strains when grown at 37°C for 6 h. Error bars show mean ± SEM. Cartoons represent nuclear morphology (black) in large-budded cells (gray). *n*, a minimum number of cells analyzed, N=4. Statistical analysis was done by two-way ANOVA using Tukey’s multiple comparisons test (**p=0.0056, ***p=0.0006). (B) Overnight grown cells of wild-type, single and double mutants were 10-fold serially diluted and spotted on YPD at 30°C and 37°C and in the presence of TBZ (50 µg mL^-1^) at 30°C. Plates were photographed after 48 h of incubation. (C) Schematic showing steps involved in the arrest and release of wild-type and *atg11*Δ cells to study Pds1 protein dynamics after metaphase arrest. Western blot analysis shows the expression of Pds1-6xHA in wild-type and *atg11*Δ cells released in YPD at 30°C after G1 and metaphase arrest using α-factor and nocodazole respectively. Cells were collected every 20 min to prepare protein samples and to quantify the budding index (*top*). Protein levels of GAPDH were used as a loading control. Pds1 normalized values are indicated below each lane. The experiments were repeated twice with similar results. (D) A bar diagram representing the proportion of cells with bioriented (light blue) or mono-oriented kinetochores (*CEN3*-GFP) (dark gray) in each strain grown at 37°C. *n* represents the minimum number of large-budded cells (budding index of >0.6) analyzed in three independent biological replicates. Wild-type (*n*>101), *atg11*Δ (*n*>106). Error bars show mean ± SEM. Statistical analysis was done using two-way ANOVA for multiple comparisons (****p<0.0001). (E) Time-lapse images showing Cdc14 (Cdc14-GFP) and spindle (Tub1-mCherry) dynamics during the cell cycle in wild-type (*top*) and *atg11*Δ (*bottom*) cells after G1 arrest and release at 37°C. The yellow arrowheads show Cdc14 release from the nucleolus. Scale bar, 5 μm. (F) The violin plot displays the time duration for wild-type and *atg11*Δ cells to release Cdc14 from the nucleolus, after G1 arrest and release at 37°C (*top*) or 30°C (*bottom*). Error bars show mean ± SEM. Statistical analysis was done using an unpaired *t*-test with Welch’s correction (*p=0.0108, **p=0.0014).

Cells of *atg11*Δ *mad2*Δ double mutant grew slower at 37°C and in the presence of TBZ (**Fig. 4B**). The SAC activation leads to the inhibition of the APC/C-Cdc20 complex, stabilizing securin (Pds1), thus preventing the activation of separase. Inactivation of separase impedes sister chromatid separation and therefore blocks metaphase-to-anaphase transition (*50*). Consistent with our observation of SAC-mediated defects, *atg11Δ* cells displayed stabilized Pds1 levels both upon release from nocodazole treatment and G1 arrest (**Fig. 4C** and **fig. S4A**). Our results, therefore, suggest that *atg11*Δ cells fail to establish correct kinetochore-MT attachments leading to a delayed anaphase onset.

**Fig S4.**
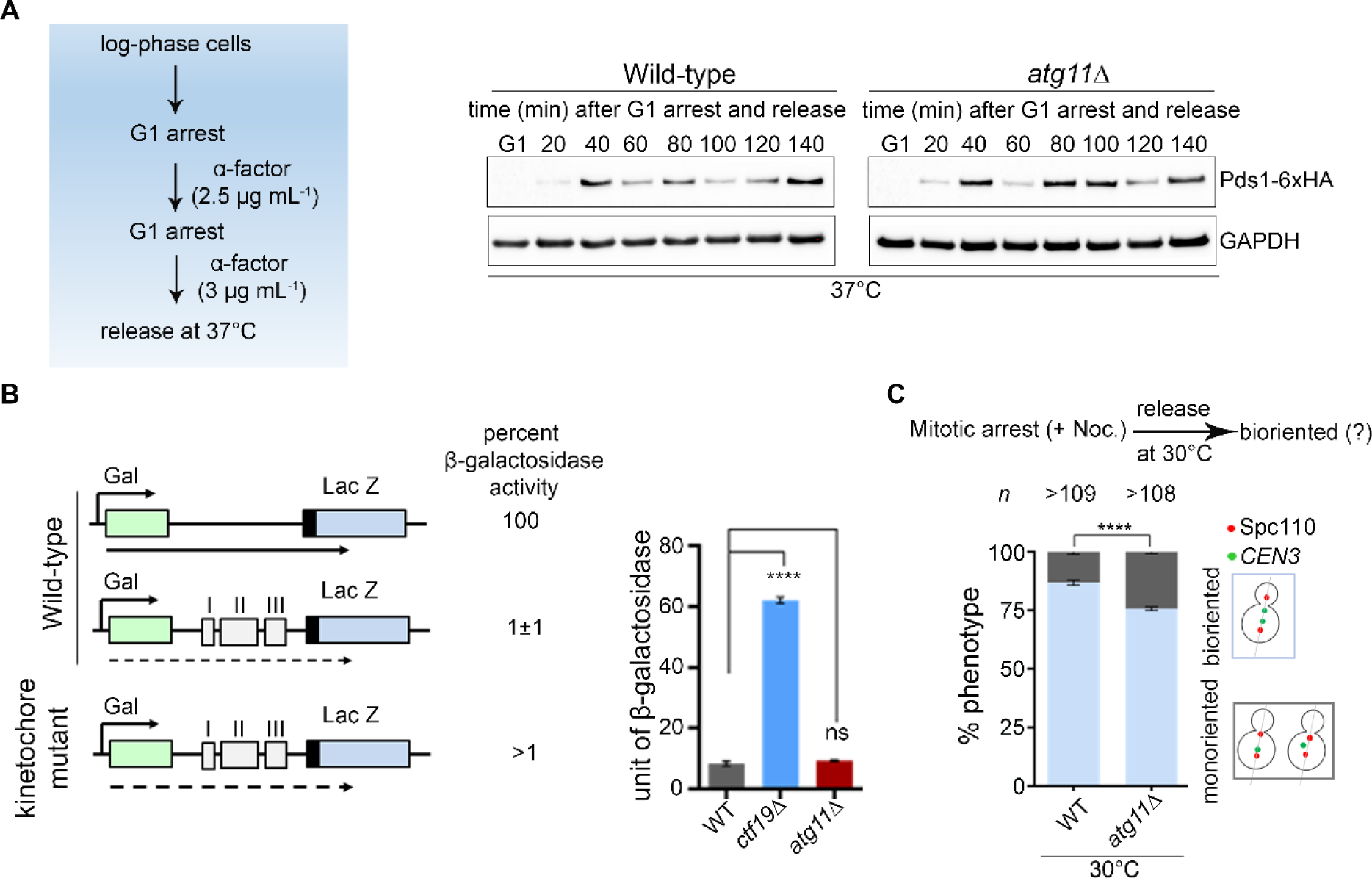
Stabilized Pds1 levels indicate defects in kinetochore-MT interactions in *atg11*Δ cells. (A) Schematic (*left*) showing steps involved in synchronizing followed by the release of wild-type and *atg11*Δ cells to study Pds1 protein dynamics after G1 arrest and release at 37°C. Western blot analysis (*right*) shows the expression of Pds1-6xHA in wild-type and *atg11*Δ cells. Protein levels of GAPDH were used as a loading control. The experiments were repeated twice with similar results. (B) Schematic (*left*) describing the transcriptional readthrough assay (*51*). Briefly, the LacZ (blue) coding sequence is in-frame with the amino-terminal actin ORF (black). *CEN* DNA (gray), labelled with I, II, and III represent CDE1, CDEII, and CDEIII respectively. Transcription is under the control of the *GAL10* promoter and the solid arrow, below the line diagram, represents maximum activity, while the width of dashed arrows represents β-galactosidase activity. A bar diagram (*right*) representing β-galactosidase levels of the corresponding strains. The measurement was performed in triplicates. Statistical analysis was done by one-way ANOVA using Dunnett’s multiple comparisons test (p<0.0001). (C) A bar diagram representing the proportion of cells with bioriented (light blue) or mono-oriented kinetochores (*CEN3*-GFP) (dark gray) in each strain grown at 30°C. *n* represents the minimum number of large-budded cells (budding index of >0.6) analyzed in three independent biological replicates. Error bars show mean ± SEM. Statistical analysis was done using two-way ANOVA for multiple comparisons (****p<0.0001).

Does Atg11 help in maintaining the structural integrity of kinetochores or in regulating MT dynamics? The *CEN* transcriptional read-through assay (*51*) ruled out the role of Atg11 in the maintenance of the kinetochore ensemble (**fig. S4B**). The kinetochore and SAC mutants display growth defects in the presence of MT poisons due to a defect in chromosome segregation. An increased chromosome mis-segregation upon loss of Atg11 and the metaphase arrest in the presence of TBZ in *atg11*Δ cells suggest that the accuracy of chromosome segregation is compromised. To test this, we monitored the fidelity of chromosome segregation in a strain with *CEN3* labeled with LacO repeats and LacI tagged with GFP (*52*). Most wild-type cells displayed bioriented sister kinetochores, as studied by co-localizing Spc110-mCherry with *CEN3*-GFP, after recovery following nocodazole treatment. However, a significant increase in mono-oriented kinetochores was observed in *atg11*Δ cells (**Fig. 4D** and **fig. S4C**). Taken together, our results reveal that Atg11 plays a critical role in cell cycle progression in conditions that destabilize MTs either by nocodazole treatment or by growing cells at 37°C. The frequency of erroneous kinetochore-MT attachments increases in the absence of Atg11 under such conditions.

As cells progress from metaphase to anaphase, the serine/threonine phosphatase, Cdc14, is first released from the nucleolus into the nucleus by the Cdc fourteen <u>e</u>arly <u>a</u>naphase release (FEAR) network in yeast. Cdc14 release is required to inactivate cyclin-dependent kinases (CDKs) during exit from mitosis (*53–56*). While securin (Pds1) and nucleolar protein Fob1 negatively regulate the FEAR network, it is positively regulated by separase Esp1, the kinetochore/spindle protein Slk19, Spo12, Bns1, and the polo kinase Cdc5 (*54, 57–60*). Since Pds1 levels are found to be stabilized in the absence of Atg11, we examined whether the *atg11*Δ cells display any delay in Cdc14 release. We probed for Cdc14 (Cdc14-GFP) dynamics in alpha-factor synchronized G1 cells. Indeed, *atg11*Δ cells showed a significant delay in Cdc14 release from the nucleolus (**Fig. 4E and F**).

The FEAR pathway regulates multiple anaphase events, including timely exit from mitosis, midzone spindle stabilization, segregation of ribosomal DNA, and nuclear positioning (*61*). Separase and Slk19 regulate the stabilization of fragile spindle midzone during anaphase by targeting the Ipl1-Sli15-Bir1 complex to the spindle midzone in early anaphase, which in turn facilitates the recruitment of the spindle-stabilizing protein Slk19 (*62–64*). The absence of either separase or Slk19 leads to the collapse of the anaphase spindle. We observed microtubule buckling in *atg11*Δ cells at anaphase, although in a small proportion of cells (5% of the live-cell) (**Fig. 5A**).

**Fig 5.**
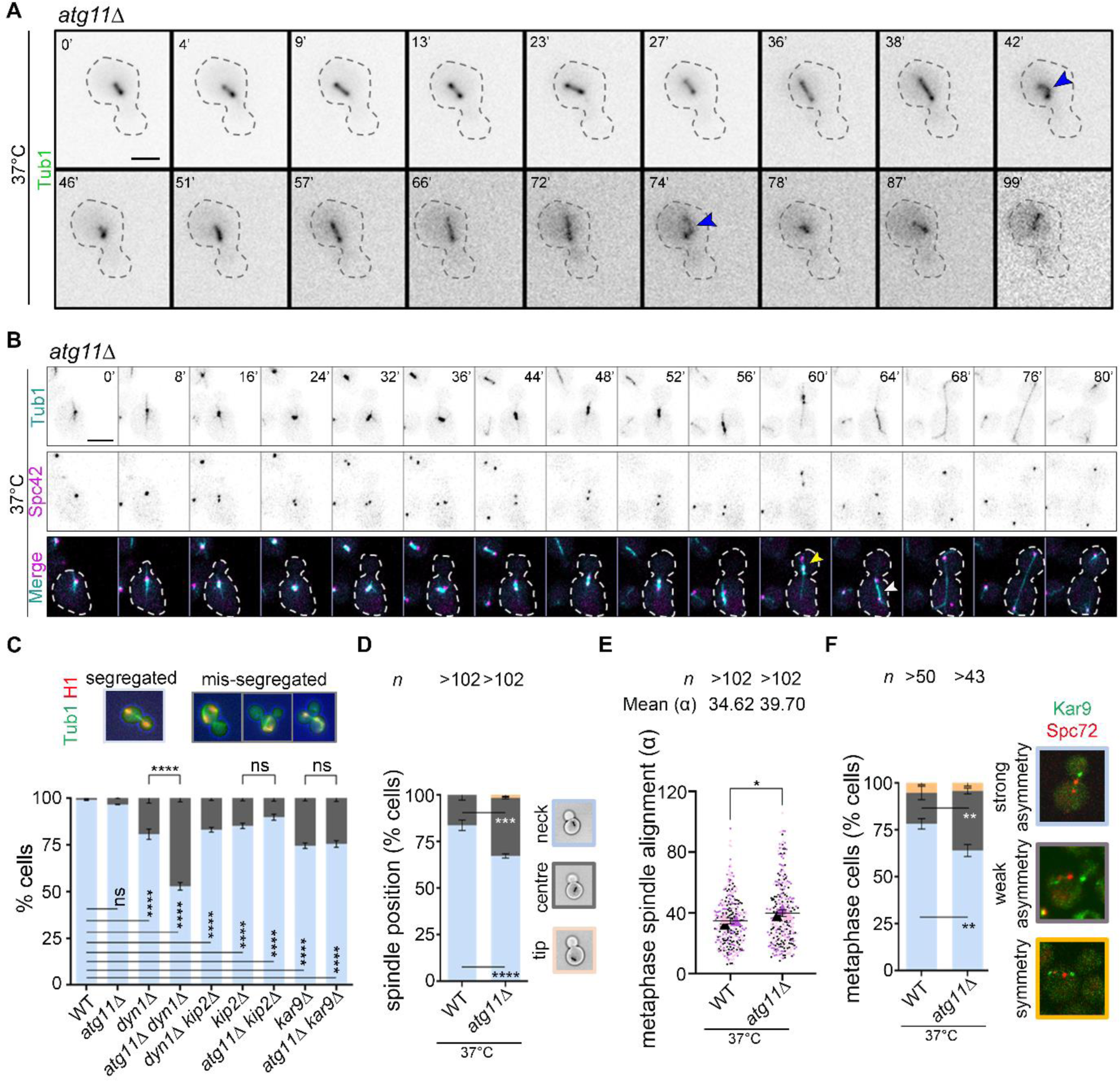
The *atg11*Δ cells exhibit Kar9-dependent spindle positioning and alignment defects. (A) Time-lapse images showing spindle (GFP-Tub1) buckling defects (blue arrowheads) at 37°C. (B) Time-lapse images showing the mitotic spindle (GFP-Tub1) and SPB (Spc42-mCherry) dynamics during the cell cycle in *atg11*Δ cells after G1 arrest followed by release at 37°C. Yellow and white arrowheads mark the movement of the mitotic spindle completely into the daughter cell and the complete elongation of the anaphase spindle in the mother cell, respectively. Scale bar, 5 μm. (C) A bar diagram showing the proportion of cells with properly segregated nuclei (light blue) or with improperly segregated nuclei represented by binucleated or multinucleated cells (dark gray). Error bars show mean ± standard error of mean (SEM). Statistical analysis was done using two-way ANOVA using Tukey’s multiple comparisons test (****p<0.0001). (D) The position of the short bipolar spindle in metaphase cells of wild-type (WT) and *atg11*Δ grown at 37°C were analyzed. *n*, a minimum number of cells analyzed, N=3. Statistical analysis was done by two-way ANOVA using Sidak’s multiple comparison test (***p=0.0001 and ****p<0.0001). (E) The angle of alignment along the mother-bud axis in metaphase cells of wild-type (WT) and *atg11*Δ grown at 37°C and displaying a short bipolar spindle were analyzed. *n*, a minimum number of cells analyzed, N=3. The statistical significance was done using an unpaired *t*-test with Welch’s correction (*p=0.0446). (F) A bar diagram showing the proportion of metaphase cells carrying Kar9-GFP either on one SPB (strong asymmetry), both the SPBs unequally (weak asymmetry) or both the SPBs equally (symmetry) in the indicated strains. Metaphase cells were identified by measuring the distance between the two SPBs (marked by Spc72-mCherry puncta). Statistical analysis was done using two-way ANOVA using Sidak’s multiple comparisons test (**p=0.0030/ 0.0050).

The FEAR network is also vital for nuclear positioning and is required to activate pulling forces at aMTs in the mother cell during anaphase. Loss-of-function of separase results in the movement of mitotic spindles into daughter cells called the ‘daughterly phenotype’ (*65*). *atg11*Δ cells displayed this phenotype in 14% of the live cells analyzed (*n*=29 live-cell) (**Fig. 5B**), along with anaphase spindle elongation in the mother cell (**Fig. 5B**, 10% of the live cells). Kar9, an MT-associated protein, regulates nuclear positioning at metaphase, is known to be dephosphorylated in early anaphase, and is suggested to be a target of Cdc14 (*12, 66*). We, therefore, examined whether Atg11 influences the Kar9-or dynein-dependent pathway of spindle positioning by examining the effect of low temperature, known to enhance the binucleated cell phenotype (*23*). The *dyn1*Δ, *kip2*Δ, and *dyn1*Δ *kip2*Δ double mutant strains exhibited a significant increase in the proportion of binucleated cells as expected (**Fig. 5C**) (*67*). Remarkably, *atg11*Δ *dyn1*Δ double mutant cells had a significantly increased proportion of binucleated cells. These cells also displayed mitotic exit with mispositioned spindles, marked by the accumulation of multinucleated and anucleate cells (**Fig. 5C**). The *kar9*Δ and *atg11*Δ *kar9*Δ mutants had more binucleated cells than those of the wild-type (**Fig. 5C**). However, the proportion of binucleated cells of the *atg11*Δ *kar9* double mutant was similar to *kar9*Δ (**Fig. 5C**), suggesting that Atg11 contributes to spindle positioning in a Kar9-dependent manner. Since Kar9 regulates spindle positioning at metaphase, we, therefore, examined whether the inactivation of Atg11 perturbed the spindle alignment and migration of metaphase spindles to the bud-neck. We measured the distance between the metaphase spindle and the bud-neck along with the angle between the spindle axis and the mother-bud axis. The *atg11*Δ cells displayed a two-fold increase in the proportion of cells having a metaphase spindle at the central region in the mother cell (**Fig. 5D**), as well as a significantly higher angle of alignment of the metaphase spindle than the wild-type (**Fig. 5E**).

Kar9 is asymmetrically localized at the daughter-bound SPB (dSPB), which in turn is required for correct spindle positioning in the wild-type cells. Therefore, we examined if the defects in spindle positioning upon loss of Atg11 are due to alterations in the asymmetric localization of Kar9 at the dSPBs. To address this, we analyzed the distribution of Kar9, tagged with GFP, in metaphase cells co-expressing Spc72-mCherry that marks the SPB marker. While 16% of wild-type cells displayed weak asymmetric Kar9 localization, the *atg11*Δ cells displayed a two-fold increase in the proportion of cells with weak asymmetric Kar9 localization (**Fig. 5F**), thereby explaining the two-fold increase in the proportion of cells having a metaphase spindle at the central region in the mother cell upon loss of Atg11 described above. Taking these observations together, our results suggest that Atg11 facilitates timely Cdc14 release from the nucleolus during early anaphase upon timely degradation of securin, which is necessary for spindle stabilization and Kar9-dependent spindle positioning and alignment in budding yeast.

### Atg11 facilitates dynamic instability of aMTs and non-random SPB inheritance

Dynamic instability of MTs also facilitates the interaction of MTs with the bud cortex, crucial for spindle orientation and positioning in budding yeast. Various factors like motor proteins, MT-binding proteins, and MT-nucleation proteins regulate MT dynamics and proper positioning of the mitotic spindle (*67–69*). We, therefore, sought to understand if Atg11 additionally contributes to the dynamic instability of aMTs in budding yeast by being localized at the SPBs. We studied aMT (GFP-Tub1) dynamics in metaphase cells of wild-type and *atg11*Δ cells (**Fig. 6A and B**).

**Fig 6.**
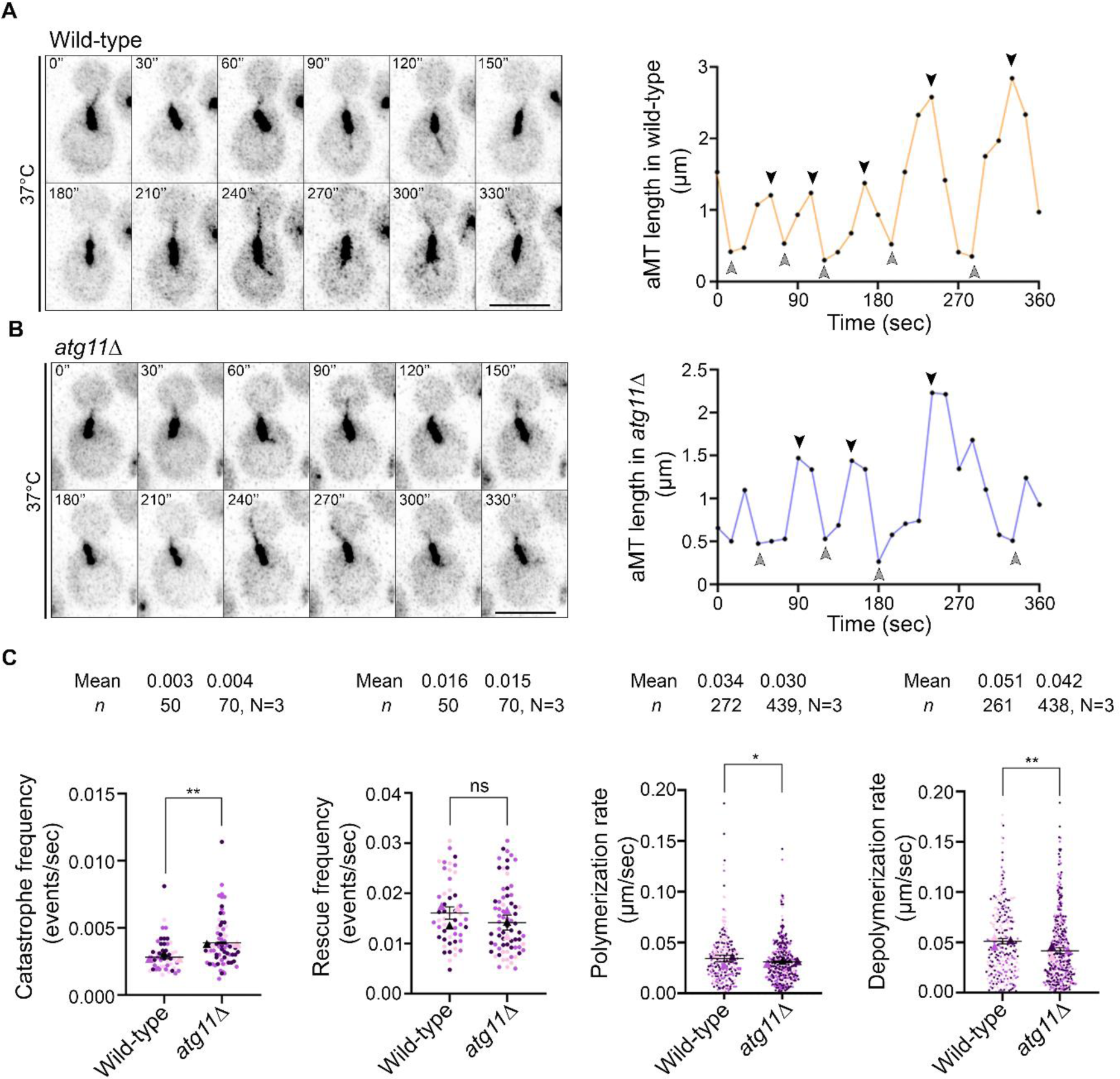
Cells lacking Atg11 have altered astral microtubule dynamics. (A) Time-lapse images (*left*) and graphs (*right*) showing the length of astral MTs (aMTs) (GFP-Tub1) in pre-anaphase cells of wild-type cells after G1 arrest and release at 37°C. (B) Similar images (*left*) and graphs (*right*) are shown for *atg11*Δ cells grown and analyzed under identical conditions as in the wild-type. Scale bar, 5 μm. Black and gray arrowheads represent the catastrophe and rescue events, respectively. (C) Scatter plot displaying catastrophe frequency, rescue frequency, polymerization rate, and depolymerization rate in wild-type and *atg11*Δ cells as indicated. Error bars show mean ± standard error of mean (SEM). Statistical analysis was done using an unpaired *t*-test with Welch’s correction (**p=0.0022/0.0015, *p=0.0443).

The *atg11*Δ cells displayed a significantly increased catastrophe frequency (**Fig. 6C**), while the rescue frequencies were comparable to those of the wild-type (**Fig. 6C**). A closer examination revealed a significant increase in the number of catastrophe events, decreased aMT length at catastrophe and lower time spent in polymerization before catastrophe in *atg11*Δ cells (**fig. S5A, B and C**).

**Fig S5.**
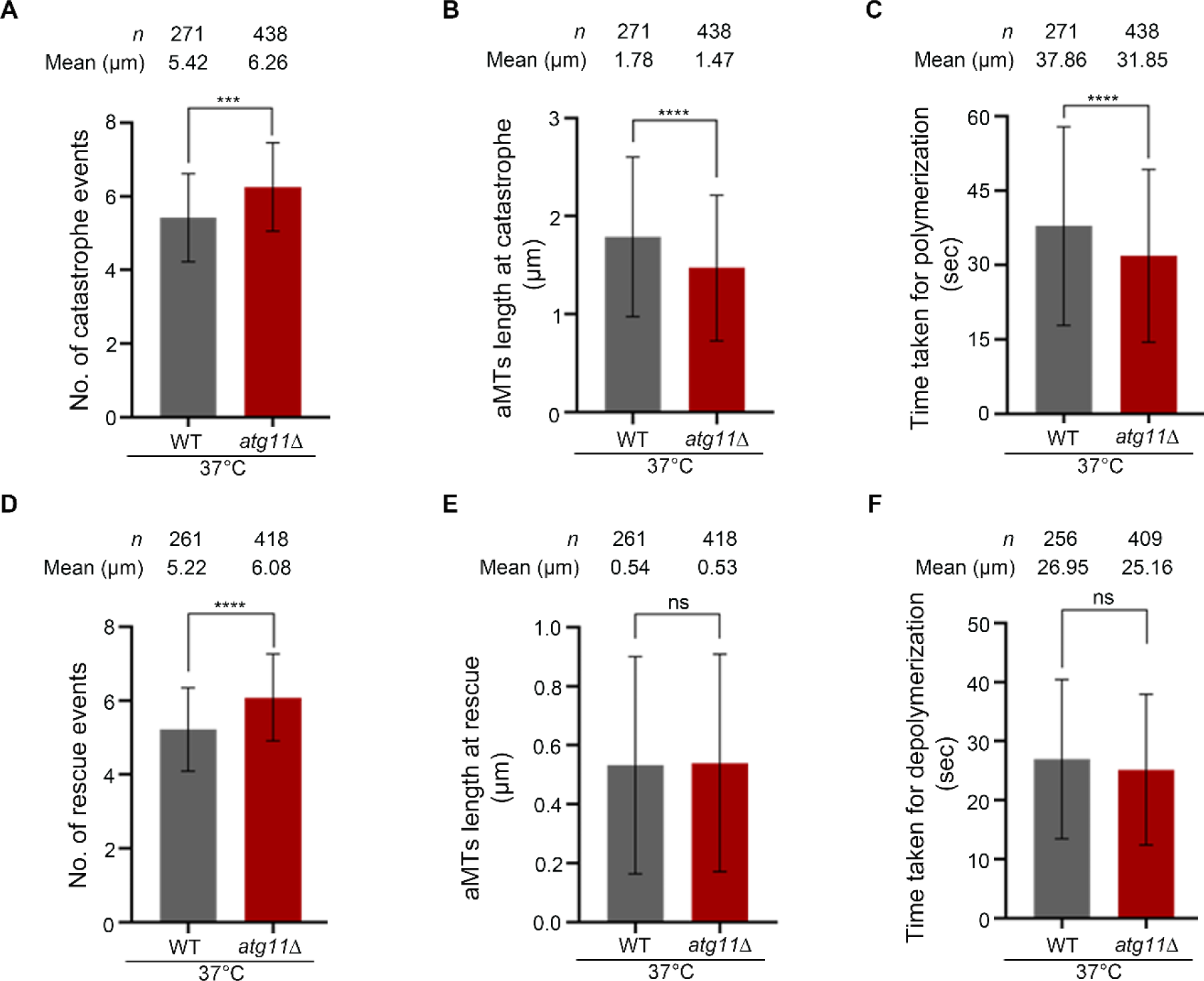
The *atg11*Δ cells display an increased number of catastrophe and rescue events, a shorter aMT length at catastrophe and a shorter time span for polymerization. (A) Bar diagram showing the number of catastrophe events in wild-type and *atg11*Δ cells. (B) Bar diagram displaying aMT length at catastrophe. (C) Bar diagram representing the time taken for polymerization before the catastrophe event. (D) Bar diagram showing the number of rescue events. (E) Bar diagram displaying aMT length at rescue. (F) Bar diagram representing the time taken for depolymerization before the rescue event. Error bars show mean ± SD. Statistical analysis was done using an unpaired *t*-test with Welch’s correction (***p=0.0003, ****p<0.0001).

The *atg11*Δ cells also showed a significant increase in the number of rescue events (f**ig. S5D**). However, there was no difference observed in the minimum length of aMTs and the time spent for depolymerization before the rescue onset between wild-type and *atg11*Δ cells (**fig. S5E and F**). Remarkably, loss of Atg11 lowered both polymerization and depolymerization rates of aMTs in *atg11*Δ cells (**Fig. 6C**). Consequently, the length of aMTs in *atg11*Δ cells was also significantly shorter at metaphase and anaphase stages of the cell cycle (**fig. S6A, B and C**). Taken together, these results revealed that Atg11 facilitates dynamic instability of aMTs and prevents catastrophe frequency in budding yeast.

**Fig S6.**
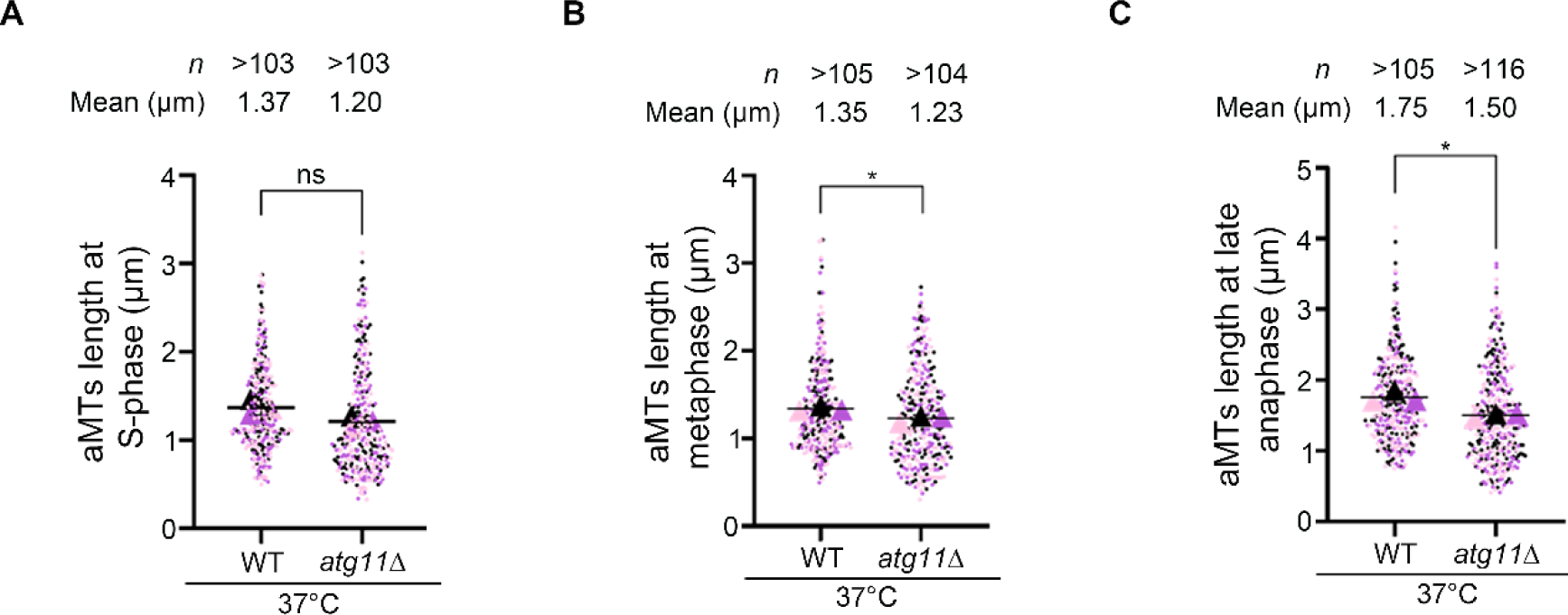
The *atg11*Δ cells exhibit shorter aMTs. (A) Scatter plot displaying aMT length at S-phase (<1 µm mitotic spindle) in wild-type (WT) and *atg11*Δ cells grown at 37°C. (B) Scatter plot representing aMT length at metaphase (1.5-2 µm mitotic spindle) in wild-type (WT) and *atg11*Δ cells grown at 37°C. (C) Scatter plot showing aMT length at late anaphase (>7 µm mitotic spindle) in wild-type (WT) and *atg11*Δ cells grown at 37°C. *n*, a minimum number of cells analyzed, N=3. The statistical significance was done using an unpaired *t*-test with Welch’s correction (*p=0.0194/0.0191).

Both asymmetric localization of Kar9 at the dSPB and proper integrity of aMTs ensure the asymmetric inheritance of the old SPB (dSPB) into the daughter bud (*12, 16*). With a random Kar9 distribution during metaphase and shorter aMTs due to loss of Atg11, we examined whether *atg11*Δ cells display any defects in SPB inheritance. To test this possibility, we tagged Spc42 (SPB marker) with mCherry. The slow folding kinetics of mCherry and ordered assembly of SPBs allow age discrimination with the new SPB (mSPB) being significantly dimmer than the dSPB (*31*). Deletion of *KAR9*, known to display random SPB inheritance (*16*), led to a randomized SPB inheritance relative to the wild-type (**Fig. 7A**).

**Fig 7.**
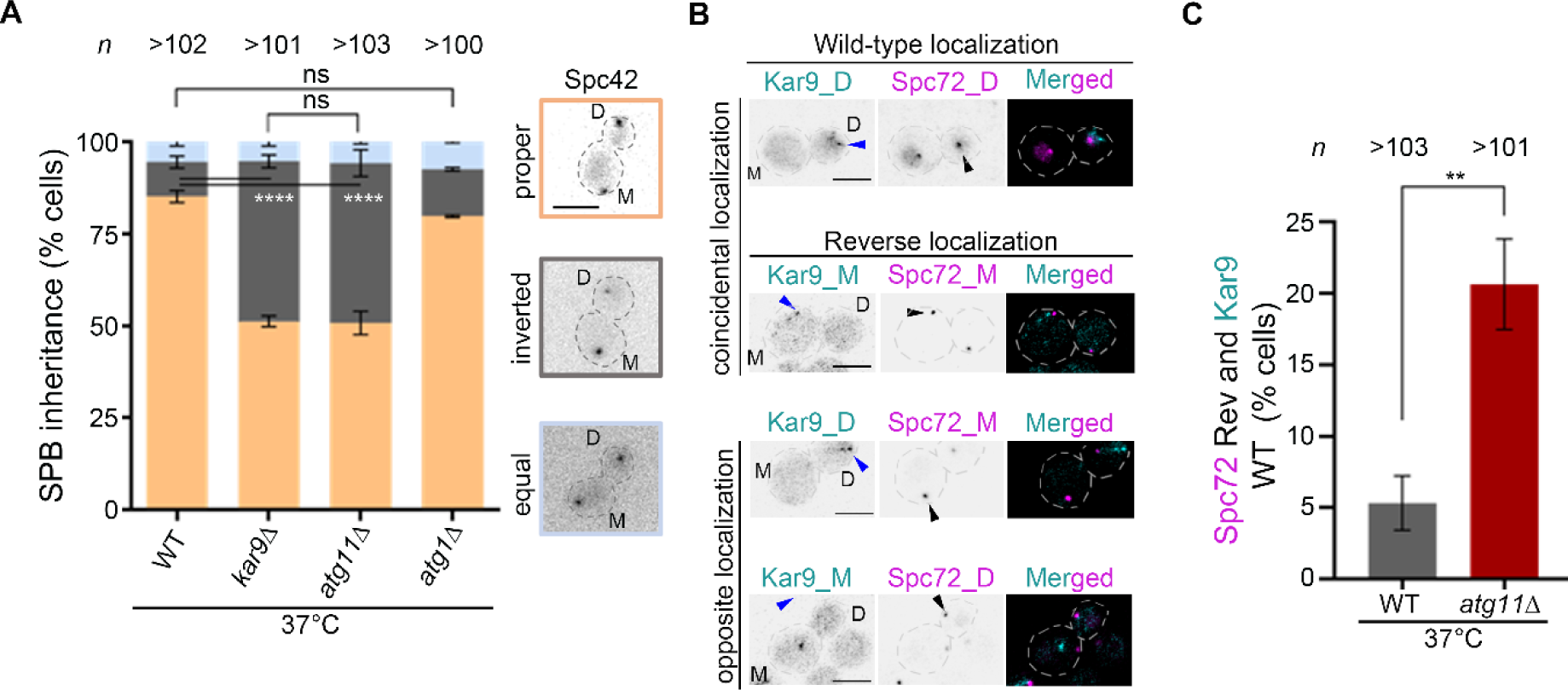
Spc72-mediated SPB inheritance is significantly altered in *atg11*Δ cells. (A) A bar diagram displaying the proportion of cells having proper (dSPB in the daughter cell), reversed (dSPB in the mother cell), or symmetric (SPBs are indistinguishable) SPB inheritance in the indicated strains is based on the Spc42-mCherry signal intensity at 37°C. *Right*, representative images of above mentioned SPB inheritance are shown. M and D represent the mother and daughter cells respectively. *n*, a minimum number of cells analyzed, N=3. Statistical analysis was done by two-way ANOVA using Dunnett’s multiple comparisons test (****p<0.0001). Scale bar, 5 μm. (B) Representative images displaying coincidental or opposite distribution of Kar9 (cyan) and Spc72 (magenta) in wild-type (WT) and *atg11*Δ cells, exhibiting wild-type-like asymmetric and predominant accumulation of both proteins in the daughter bud (D) or reversed such that asymmetric and predominant accumulation of these proteins in the mother (M) cell. Asymmetric localization of Kar9-GFP and Spc72-mCherry is marked by blue and black arrowheads, respectively. Scale bar, 5 μm. (C) Bar diagram representing the proportion of cells displaying inverted Spc72 localization when Kar9 is asymmetrically localized in daughter cells at 37°C. *n*, a minimum number of cells analyzed, N=3. Error bars show mean ± standard error of mean (SEM). The statistical significance was done using an unpaired *t*-test with Welch’s correction (**p=0.0041).

Strikingly, *atg11*Δ cells showed a four-fold increase in SPB inheritance defects (**Fig. 7A**). Notably, autophagy mutant cells lacking a key autophagy protein Atg1 segregated SPBs like in the wild-type cells (**Fig. 7A**), hinting that SPB inheritance is autophagy-independent in budding yeast. A previous report on the localization interdependency of Spc72 and Kar9 onto the SPBs, placed Spc72 upstream of Kar9 (*70*), while a recent study demonstrates Cdc5 as a molecular timer facilitating timely recruitment of both Spc72 and Kar9 onto the SPBs (*71*). Based on the SPB inheritance defects, we asked whether Atg11 contributes to the asymmetric localization of Spc72 and Kar9 at the SPBs. Strikingly, the loss of *ATG11* randomized the SPB localization of Spc72 (**Fig. 7B and C**). Taken together, we conclude that being localized at the SPBs, Atg11 prevents catastrophe frequency of aMTs, crucial for proper aMT length and asymmetric SPB inheritance in budding yeast.

## Discussion

We unravel an unusual function of an autophagy-related protein, Atg11, in maintaining the high-fidelity transmission of genetic material through many generations. Our results demonstrate that *atg11*Δ cells show a high rate of plasmid and chromosome loss. How does an autophagy-related protein, predominantly distributed in the cytoplasm, regulate chromosome segregation? Strikingly, single-molecule tracking provides evidence of a significant and dynamic pool of Atg11 molecules associated with SPBs. Loss of Atg11 leads to the accumulation of a large proportion of cells with mono-oriented chromosomes and SAC-mediated delay of cell cycle progression. This is supported by the stability of securin protein Pds1 observed in *atg11*Δ mutant cells. *atg11*Δ cells show the appearance of FEAR-defective phenotypes of spindle stabilization and loss of Kar9-dependent nuclear positioning. In addition, aMT catastrophe is significantly altered in *atg11*Δ mutant cells. This is evidenced by a higher proportion of cells with short aMTs. Without Atg11, many cells lost asymmetric localization of both Spc72 and Kar9, leading to the loss of asymmetric SPB inheritance. We propose that by localizing at SPBs, Atg11 regulates the biased distribution of factors between daughter and mother SPBs. This impacts MT dynamics, both in the nucleus and cytoplasm and a set of coordinated events that ensure precise chromosome segregation (**Fig. 8**). The non-canonical role of Atg11 in maintaining ploidy is significant, especially in light of an increasing body of evidence that suggests defects in MT-mediated cellular processes generate aneuploidy, a cellular state commonly observed in diseases such as cancer (*72–77*).

**Fig 8.**
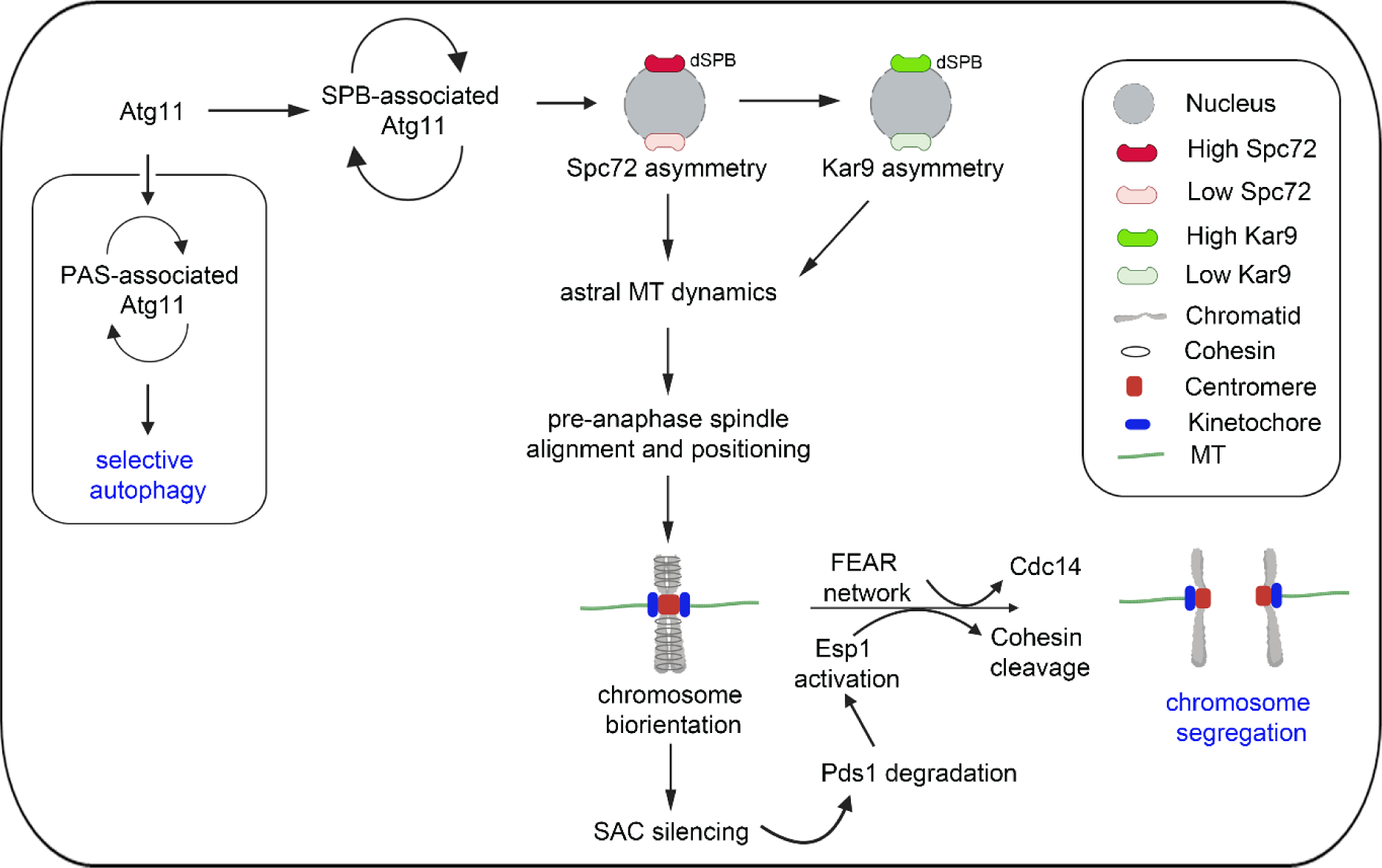
SPB-associated Atg11 ensures high-fidelity chromosome segregation in budding yeast. This study reveals two distinct pools of Atg11 in budding yeast. A lower proportion of Atg11 which is less dynamic canonically localizes at the pre-autophagosomal structures (PAS) onto the vacuolar membrane (*48*), necessary for selective autophagy. While the more dynamic Atg11 localizes at the SPBs in a higher proportion which is critical for high-fidelity chromosome segregation. The asymmetric Kar9 localization together with the dynamic instability of MTs is crucial for pre-anaphase spindle positioning and alignment, and chromosome biorientation. Once biorientation is achieved, SAC is silenced, Pds1 gets degraded, separase is activated, and Cdc14 is released from the nucleolus via the FEAR pathway leading to proper sister chromatid separation (*78*). Atg11 ensures asymmetric localization of both Spc72 and Kar9 at the dSPBs, critical for non-random SPB inheritance.

Based on the single-molecule tracking (SMT) assay, our study validates two distinct pools of Atg11 in the budding yeast, including a less dynamic cytoplasmic pool of Atg11 required for selective autophagy. Consisting of four coiled-coil domains, Atg11 functions as a scaffolding protein, with the fourth coiled-coil domain (CC4) of Atg11 important for selective autophagy (*48*). The second pool of Atg11, being more dynamic, uncovers a non-canonical localization of Atg11 to SPBs. The interaction of Atg11 with Spc72 at SPBs suggests a repurposing of the scaffolding function of Atg11 in budding yeast, which undergoes closed mitosis. We posit that being a bridging protein, Atg11 likely interacts, recruits, or positions proteins key to the dynamic instability of MTs and asymmetric distribution of Spc72. The polo-like kinase, Cdc5, facilitates the asymmetric localization of Spc72 and Kar9 at the dSPBs crucial for asymmetric SPB inheritance (*71*). Cdc5 also displays a physical association with Spc72 (*79*) and safeguards SPB asymmetry. Our study suggests the loss of non-random SPB inheritance in the absence of Atg11. Therefore, it is tempting to speculate that Atg11 and Cdc5 may interact *in vivo* and such an interaction facilitates asymmetric localization of Spc72 and Kar9. It remains to be seen if Atg11 brings Cdc5 to dSPB or *vice-versa*.

The alignment of the mitotic spindle along the polarity axis ensures proper inheritance of DNA between the two cells during asymmetric cell division. The asymmetric localization of Kar9 at the dSPBs together with long and dynamic aMTs is crucial for the positioning and alignment of the pre-anaphase spindle. Consistent with this, a loss of function of proteins affecting either aMT dynamics or positioning of pre-anaphase spindle displays misaligned and mispositioned mitotic spindle (*13, 17, 68, 69, 80, 81*). A larger pool of Atg11 molecules around SPBs may explain how Atg11 regulates MT dynamics primarily at the SPBs. Indeed, we show that Atg11 plays a role in regulating aMT instability, maintains overall aMT length, and prevents an increased catastrophe frequency (**Fig. 8**). Whether Atg11 positively regulates proteins like Kip2 and Stu2 (*82*), which prevent the catastrophe frequency, or it negatively regulates proteins like Kip3 (the Kinesin-8 or Kinesin-13 family) or Kar3 which destabilizes MTs *in vivo* (*26, 82, 83*), remains to be studied.

Dynamic instability, wherein MTs exhibit cycles of growth and shrinkage is central to high-fidelity chromosome segregation and is therefore tightly regulated during the cell cycle by several proteins and post-translational modifications (*84*). Dynamic MTs facilitate the timely attachment of MTs with the kinetochores in the nucleus and are crucial for the generation of tension between sister chromatids during the cell cycle (*68*). In this study, we demonstrate the significance of Atg11 in maintaining MT integrity, especially under conditions that perturb their rate of polymerization-depolymerization. The presence of Atg11 silences the SAC by facilitating chromosome biorientation, leading to securin degradation and timely Cdc14 release by activating the FEAR pathway (**Fig. 8**). However, it is unclear how being a cytosolic protein, Atg11 facilitates proper kinetochore-MT attachments within the nucleus during closed mitosis in *S. cerevisiae*. In budding yeast, several proteins regulate MT dynamics both in the nucleus and cytosol. Localized to SPBs, spindles and the plus-end of aMTs, the MT-binding protein Stu2 is known to regulate kinetochore-MT attachment, aMT, and interpolar MT dynamics (*68*). Kar3, a minus-end directed motor, localizes to SPBs, mediates MT depolymerization at SPBs and a loss of Kar3 displays a SAC-dependent delay in the cell cycle (*85, 86*). Whether Atg11 regulates the dynamics of MTs via these factors or whether some molecules of Atg11 also migrate to the nucleus remains an area of further investigation. Our findings are significant in light of the autophagy-independent role of Beclin-1 in accurate kinetochore anchoring to the spindle during mitosis in cells undergoing open mitosis (*87*). Thus, like Beclin-1, Atg11 could be another example of an autophagy-related protein regulating MT dynamics essential for both kinetochore-MT interactions and spindle positioning during the cell cycle. Depending on the types of mitoses, the players might differ in performing a non-canonical function. Several lines of evidence have established an interplay between errors in the spindle alignment with aging and aneuploidy leading to the development of cancer and neurodegenerative diseases (*28–32*). Likewise, perturbations in the autophagy function also contribute to various human diseases like neurodegenerative diseases, cancer, lysosomal storage disorders, and inflammatory disorders (*88, 89*). Our work adds to the importance of cellular functions carried out by transient protein-protein interactions vital for cellular fitness under specific conditions. Therefore, dissecting these dynamic interactions essential for crosstalk between various biological processes becomes imperative.

## Materials and Methods

### Yeast strains, plasmids, and primers

All the strains used in this study are specified in Supporting Table S1. Primer sequences and the plasmids used for strain construction are specified in Supporting Table S2, and Supporting Table S3, respectively.

### Media and growth conditions

Yeast strains were grown on YPD media (yeast extract 1%, peptone 2%, and dextrose 2%) at 30°C unless stated otherwise. For the growth assay in the presence of thiabendazole (TBZ), overnight grown cultures of wild-type, autophagy mutants, *ctf19*Δ, *mcm16*Δ, and *ATG11*-complemented strains were serially diluted and spotted on YPD containing 50 µg mL^-1^ and imaged on 2 dpi or 36 h. The analyses for figures 3C and 7A-C were done by growing the cells at 37°C for 6 h.

### Interaction network

First, we constructed a protein-protein interaction map by identifying the proteins that have been reported to either genetically or physically interact with each of the Atg proteins. We further extended the network to include the proteins that are known to interact with any of the primary interactor proteins. We gleaned these connections using Perl scripts to query the latest curation of the BioGRID yeast interactomes (*43*). To this end, only those protein-protein interactions were retained which were confirmed and corroborated across three or more independent studies.

### Construction of yeast strains

The *ATG11* gene was deleted from the *S. cerevisiae* genome by one-step gene replacement (*90*). A 40-bp homology region corresponding to the upstream and downstream sequence of *ATG11* having *HPH* marker providing resistance against hygromycin B was amplified from pAG32 using HR34 and HR35 primer pairs. Similarly, *ATG11* was deleted in F2351 (*31*), with Kar9-GFP and Spc72-mCherry tagged, using primers (HR34 and HR35) carrying 40-bp homology regions from upstream and downstream of the *ATG11* gene that were used to amplify the *HPH* marker sequence from pAG32. The *ATG11* gene deletion in the desired transformants was confirmed by PCR using 5’ and 3’ junction primers (HR36 and HR37).

Double deletion mutants were obtained either by transforming the above *ATG11* deletion cassette in a single deletion mutant background or by crossing a single deletion mutant in *MAT***a** background having the G418 marker with *atg11*Δ cells in *MATα* (BY4742) background having the *HPH* marker. The diploid cells were sporulated and after digestion of the cell wall and vortexing, the cells were plated first on YPD with hygromycin. Growing colonies were patched on G418 and colonies grown on both the selection plates were mated again with both *MAT***a** (BY4741) and *MATα* (BY4742) to differentiate between haploid and diploid cells. Cells showing shmoo formation with only one mating type were taken forward for the assays.

For *atg11*Δ complementation, a ∼3.9 kb fragment comprising the upstream and full-length sequence of *ATG11* was fused with a ∼800 bp of *HIS3* fragment with its promoter having a 40-bp homology sequence of *ATG11* at its 3’ end. The purified PCR product was transformed in the *atg11*Δ cells and the desired transformants were confirmed using HR116 and HR117 primers.

### Construction of a strain expressing GFP-Tub1 and Spc42-mCherry under its native promoter

To study the dynamics of MTs, *TUB1* was tagged with GFP using the plasmid, pAFS125 (*91*). To visualize SPBs, Spc42, a component of the central plaque of SPB, was tagged with mCherry using the primers HR142 and HR143.

### Construction of *MAD2* deletion strain

The *MAD2* gene was deleted by one-step gene replacement (*90*), having a 40-bp homology region each from upstream and downstream of the gene using HR52 and HR53 primer pairs. These primers were used to amplify the *HPH* marker from pAG32. The desired deletion strain was confirmed by PCR using 5’ and 3’ junction primers (HR66 and HR67).

### Construction of Pds1-6xHA tagged strain

A PCR fragment containing 6xHA and the *HPH* selection marker was amplified from pYM16 (*92*) using the primer pair RA140 and RA141. To obtain Pds1-6xHA, the PCR product was transformed in wild-type BY4741 cells generating the strain ScRA403. Transformants were selected using the hygromycin selection marker. To tag Pds1 in *atg11*Δ, *MAT***a** RA403 was crossed with MHR08 (*MAT*α). Diploids were selected using methionine and lysine prototrophy. Diploids were sporulated and haploid progeny were identified by mating with both *MAT***a** and *MATα* strains. The desired strain (ScRA405) was confirmed for both *ATG11* deletion and Pds1 tagging using the primer sets HR36 and HR37 and RA146 and RA147, respectively. Pds1-6xHA tagging both in wild-type and *atg11*Δ cells were confirmed by western blot analysis.

### Construction of strains for Bimolecular Fluorescence Complementation (BiFC) assay

We used N-terminal protein tagging vectors, pFA6a-KanMx6-p*GAL1*-VN173 and pFA6a-His3Mx6-p*GAL1*-VC155 for tagging either Atg11 or Spc72 (*46*). VN and VC designate N-terminal and C-terminal fragments of Venus, a variant of yellow fluorescent protein, respectively (*46*). VN173-Atg11 and VC155-Atg11 were amplified using HR296 and HR297 and HR296 and HR298 primer pairs respectively. Similarly, VN173-Spc72 and VC155-Spc72 were amplified using HR299 and HR300 and HR299 and HR301 primer pairs respectively. VN-tagged fragments with the G418 selection and VC-tagged fragments with the *HIS3* selection were transformed into BY4741 and BY4742 backgrounds respectively. The haploid colonies obtained were mated and selected on G418 plates without histidine to obtain diploid cells expressing both fragments of the fluorophore. The control diploid cells expressing either VN-Spc72 or VC-Atg11 co-expressing Spc42-mCherry were generated by mating BY4741 expressing VN-Spc72 with BY4742 or BY4742 expressing VC-Atg11 with BY4741.

### Construction of strains for Single Molecule Tracking (SMT) assay

The *PDR5* gene, coding for the membrane transporter protein Pdr5 was deleted to allow the retention of the HaloTag ligands (JF646) inside the yeast cell (*93*). The *PDR5* gene was deleted in the MHR90 *MAT***a** strain (sfGFP-Atg11 Spc42-mCherry) using a 40-bp homology region each from upstream and downstream of the *PDR5* gene using HR333 and HR334 primer pairs. These primers were used to amplify the *HPH* marker from pAG32. The desired deletion strain was confirmed by PCR using 5’ and 3’ junction primers (HR335 and HR336). For labelling Atg11 using HaloTag Ligand (HTL), the *ATG11* gene was fused with *-*HaloTag using the *URA3* selection marker by homologous recombination. The HaloTag cassette was amplified from pTSK561 (Addgene: 190816) using HR329 and HR330 primer pairs and transformed in yTK1501 *MAT*α strain. The tagged strains were confirmed by HR36 and HR37 primer pairs. The diploid strain, expressing both sfGP-Atg11 and Atg11-HaloTag, was created by yeast mating and the diploid cells were selected on CM-ura media.

### Stability of monocentric plasmid

The *atg1*Δ, *atg6*Δ, *atg11*Δ, *atg15*Δ, *atg17*Δ, *ctf19*Δ, and the isogenic wild-type strains were transformed with pRS313 (*94*), a centromeric plasmid, and transformants were selected on a dropout media lacking histidine (SD-his). The transformants were then grown in a non-selective media, YPD, for 7-10 generations, followed by plating to obtain single colonies. The single colonies were subsequently patched onto SD-his (selective) and YPD (non-selective) media. The mitotic stability was calculated by the number of colonies that were able to grow on selective media (SD-his) per 100 colonies grown on non-selective media (YPD).

### Chromosome loss assay

The chromosome loss assay was performed in a strain harboring an extra non-essential chromosome that carries *SUP11* that suppresses *ade2* mutation in the assay strain, YMH58a as previously described (*95*). Cells retaining this extra chromosome grow as white-colored colonies, complementing the *ade2* mutation. Loss of extra chromosome leads to the red-pigmented colony, displaying *ade2* mutant phenotype, implying loss of *SUP11*. A single white colony of wild-type, *atg11*Δ, and *ctf19*Δ cells were grown in non-selective media (YPD) for 5-7 generations, both at 30°C and 37°C. The cells were plated on YPD. The number of colonies showing chromosome loss at the first division was scored for each condition in three independent biological experiments.

### *In vivo* assay for kinetochore integrity: *CEN* transcriptional read-through assay

The *CEN* transcription read-through assay was done as previously described (*51*). Briefly, the reporter plasmid, pAKD06+*CEN4* contains a β-galactosidase reporter gene with the selectable marker *URA3* was transformed in wild-type, *ctf19*Δ and *atg11*Δ cells and selected on SD-ura media. The single colony was inoculated in SD-ura media, containing 2% galactose and 0.3% raffinose and was grown to 1 OD_600_. The cell pellet was washed with water and resuspended in 1 mL of Z-buffer (60 mM Na_2_HPO_4_, 40 mM NaH_2_PO_4_.2H_2_O, 10 mM KCl, 5 mM β-ME). 0.1 mL of this was taken for the determination of OD_610_. 0.1 mL of Z-buffer was added to the remaining 0.9 mL of cells. The cells were then permeabilized by adding 50 µL of 0.1% SDS and 100 µL of chloroform (incubated at 30°C for 15 min). 0.2 mL of 4 mg mL^-1^ ONPG (ortho-nitrophenyl-β-galactoside) was added to the cell suspension and incubated at 30°C till yellow colour developed. The reaction was stopped by adding 0.5 mL of 1 M Na_2_CO_3_. Cells were spun down at 10,000 *g* and the clear supernatant was transferred to a fresh tube. The optical density of this solution was measured at 420 and 550 nm. The enzyme activity was normalized with respect to cell density. Units of β galactosidase were measured as:

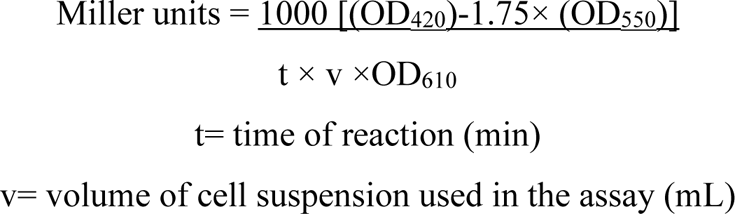

### Live-cell imaging

For tethering the cells to the glass-bottom dishes for live-cell imaging, the dishes were treated with 6% Concanavalin A (Catalogue no. C2010, Sigma-Aldrich) for 10 min. Wild-type and *atg11*Δ cells carrying GFP-Tub1 (spindle) and Spc42-mCherry (SPBs), or Cdc14-GFP and Tub1-mCherry were grown overnight in CM medium, re-inoculated in CM at 0.2 OD_600_ and grown for 3 h. The cells were first arrested at G1 using 3.0 µg mL^-1^ α-factor at 30°C (Merck, Cat. No. T6901). After 90 min, 3.0 µg mL^-1^ α-factor was again added and incubated for another 90 min. Cells were washed thrice with CM medium and released into Concanavalin A-treated glass bottom dishes and incubated at 30°C or 37°C for 10 min. Live-cell imaging was performed on an inverted confocal microscope (ZEISS, LSM880) equipped with a temperature-controlled chamber (Pecon incubator, XL multiSL), a Plan Apochromat 63x NA oil 1.4 objective and GaAsP photodetectors. For time-lapse microscopy, images were captured at a 4 min interval with 5% intensity exposure with 0.5 µm Z-steps using 488 nm and 587 nm for GFP and mCherry as excitation wavelength. All the images displayed were maximum intensity projections of images for each time using ImageJ.

### Microscopic image acquisition and processing

Wild-type and deletion mutants carrying either GFP-Tub1 (spindle) or Spc42-mCherry (SPB marker) were grown overnight in YPD medium and re-inoculated in YPD at 0.2 OD_600_ for 3 h at 30°C. The cells were pelleted at 3,000 *g* and mounted on a glass slide having a 2% agarose pad supplemented with 2% dextrose. The pad was sealed with a coverslip. The cells were imaged using an inverted microscope (ZEISS, Axio Observer.Z1/7), a Plan-Apochromat 100x/1.40 oil DIC M27. The filters used were GFP/FITC 488, mCherry 561 for excitation, GFP/FITC 500/550 bandpass or GFP 495/500 and mCherry 565/650 or mCherry 580/750 bandpass for emission. Z-stacks at an interval of 0.5 µm were taken and maximum intensity projection for each image using ImageJ was projected.

For Figures 3A, 4E, 5A-B, 5F, 6A-B, and 7B-C, supporting figures S2B-F, S3A, S4C, and S6A-C were imaged using an inverted confocal microscope (ZEISS, LSM880) equipped with an Airyscan module, 63x Plan Apochromat 1.4 NA. 488/516 and 561/595 nm excitation/emission wavelengths were used for GFP and mCherry respectively.

### Biorientation assay

For the *CEN3* biorientation assay, overnight grown wild-type and *atg11*Δ cells were reinoculated to 0.2 OD_600_ and grown for 3 h. The cells were arrested at metaphase in the presence of 15 µg mL^-1^ of nocodazole for 2 h. The cells were then released in YPD in the presence of 50 µM bathophenanthrolinedisulfonic acid disodium salt (BCS) at 30°C or 37°C for 1 h. Cells with two *CEN3*-GFP puncta between the two SPBs (Spc110-mCherry) were considered bioriented, while those with one *CEN3*-GFP punctum closer to either of the SPBs or away from the polarity axis were considered mono-oriented. The large-budded cells with a budding index of >0.6 were used for the analysis.

### Western blot for Pds1 dynamics after nocodazole arrest

Overnight grown cells were diluted to 0.1 OD_600_ for wild-type and 0.15 for *atg11*Δ cells in 25 mL complete media. The cultures were then grown at 30°C until 0.3-0.4 OD_600_. The cells were first arrested at G1 using 2.5 µg mL^-1^ α-factor grown at 25°C (Cat. No. T6901, Merck). After 90 min, 3.0 µg mL^-1^ α-factor was again added and incubated for another 90 min. Cells were washed with water thrice and released into YPD media containing 15 µg mL^-1^ nocodazole (Cat. No. M1404, Merck) for 90 min. Metaphase-arrested cells were washed thrice in water and released in 25 mL YPD. The cultures were grown at 30°C and 1 OD_600_ equivalent culture was collected and pelleted every 20 min. The cell pellets were lysed by TCA precipitation and resuspended in 50 µL lysis buffer (0.1N NaOH, 1% SDS and 5x protein loading dye). Pds1 degradation dynamics were analyzed using a western blot. Pds1 levels were monitored using rat anti-HA antibody (Cat. No. 11867423001, Roche) and peroxidase-conjugated rabbit anti-rat secondary antibody (Cat. No. HPO14, Genei) at 1:3000 and 1:10000 dilution respectively. GAPDH levels were analyzed using mouse anti-GAPDH antibody (Cat. No. MA5-15738, Invitrogen) and HRP conjugated goat anti-mouse secondary antibody (Cat. No. ab97023, Abcam) at 1:5000 and 1:10000 dilution respectively. The blots were developed using a Chemiluminescence substrate (BioRad) and imaged using the ChemiDoc imaging system (BioRad). Pds1 levels were normalized with their respective loading control at every time point for both wild-type and *atg11*Δ cells. To trace the dynamics of Pds1, a ratio of the normalized values at each time point with its 0 min was plotted.

### GFP-Atg8 processing assay

The *atg1*Δ, *atg11*Δ, *atg17*Δ, and the isogenic wild-type strains were transformed with GFP-Atg8 pRS423, a 2 µm plasmid, and selected on a dropout media lacking histidine (SD-his). The overnight growing cells in SD-his media were reinoculated to 0.25 OD_600_ in SD-his till the OD reached 1.0. The cells were pelleted and washed once with a nitrogen starvation medium (yeast nitrogen base + 2% dextrose). 1 OD cell/mL cells were transferred to a pre-warmed nitrogen starvation medium (30°C and 37°C). 1 mL cells corresponding to 1 OD were aliquoted at 0, 2, 4, and 6 h post incubation in a nitrogen starvation medium. The cell pellets were lysed by TCA precipitation and resuspended in 50 µL lysis buffer (0.1N NaOH, 1% SDS and 5x protein loading dye). GFP-Atg8 processing assay was analyzed using a western blot. GFP-Atg8 and free GFP levels were monitored using rat anti-GFP antibody (Roche, Cat. No. 11867423001) and peroxidase-conjugated rabbit anti-rat secondary antibody (Genei, Cat. No. HPO14) at 1:6,000 and 1:10,000 dilution respectively. The blots were developed using a Chemiluminescence substrate (BioRad) and imaged using the ChemiDoc imaging system (BioRad).

### Bimolecular Fluorescence Complementation (BiFC) assay

The overnight growing cells co-expressing Spc72 fused to the N-terminal half of Venus and Atg11 tagged to the C-terminal half of the same fluorophore were reinoculated to 0.2 OD_600_ in YEP containing 2% galactose (YPG) for 3 h. To label the vacuolar membrane, FM4-64 was added at a concentration of 7.5 μM for 30 min. The cells were washed twice with 1x PBS and further incubated for 1 h in YPG before microscopy.

### Staining protocols

FM4-64 staining was performed to visualize the vacuolar membrane. Briefly, FM4-64 (Cat. No. T3166, Thermo Fischer Scientific) was added at a concentration of 7.5 μM for 30 min. The cells were washed twice with 1x PBS and further incubated for 1 h in YPG before microscopy.

Nuclei were visualized by diamidino-2-phenylindole (DAPI) (Cat. No. 62248, Thermo Fischer Scientific) staining. Briefly, the cells were fixed in 3.7% formaldehyde for 10 min. The cells were washed and treated with 0.05% Triton X-100 for 5 min. The cells were resuspended in 1x PBS and stained with 5 ng mL^-1^ DAPI for 10 min and imaged.

### Yeast two-hybrid assays

The full-length *ATG11* (∼3534 bp), *SPC72* (∼1869 bp), and *CNM67* (∼1746 bp) were PCR amplified and cloned in frame with the Gal4 activation/ DNA binding domain (AD or BD domain) in pGADC1 and pGBDC1 respectively at BamHI and SalI sites using oligos listed in Supplementary file 2. Yeast-two hybrid analyses were performed in the strain PJ69-4A as described previously (*96*). Briefly, bait and prey plasmids were co-transformed into the *S. cerevisiae* strain PJ69-4A. Positive interactions were scored by the appearance of white-coloured colonies on a synthetic defined medium containing 6 µg mL^-1^ adenine and/ or by the growth in the presence of 2 mM 3-Amino-1,2,4-triazole (3-AT) (Cat. No. A8056, Sigma-Aldrich).

### aMT dynamics

The overnight grown wild-type and *atg11*Δ cells carrying GFP-Tub1 were reinoculated to 0.2 OD_600_ and were grown for 3 h in CM-ura medium. The cells were first arrested at G1 using 3.0 µg mL^-1^ α-factor grown at 25°C (Merck, Cat. No. T6901). After 90 min, 3.0 µg mL^-1^ α-factor was again added and incubated for another 90 min. Cells were washed thrice with prewarmed CM-ura medium at 37°C and released in the same medium at 37°C till the cells reached metaphase (1.5-2.0 µm). For tethering the cells to the glass-bottom dishes for live-cell imaging, the dishes were treated with 6% Concanavalin A (Cat. No. C2010, Sigma-Aldrich) for 10 min. Cells were resuspended in CM-ura medium and were allowed to adhere for 10 min at 37°C. The unadhered cells were washed and the adhered cells were resuspended in 3 mL of CM-ura and proceeded for live-cell imaging at 37°C. The images were acquired every 15 s at 1024x1024 frame size for a total of 25 cycles. At each time point, images were taken at 0.4 µm apart, covering a total depth of 4.5-5.5 µm. Images were processed by a deconvolution process automatically performed by the Zeiss software. Image analysis was done using Fiji. As described in (*97, 98*), various events were defined as follows: (a) catastrophe frequency-the total number of polymerization-to-depolymerization transitions divided by the total time of all growth events; (b) rescue frequency-the total number of depolymerization-to-polymerization transitions divided by the total time of all shrinkage events; (c) polymerization rate-divide the net change in length by the change in time for each growth event; (d) depolymerization rate-divide the net change in length by the change in time for each shrinkage event.

### Single Molecule Tracking (SMT) assay

The diploid cells tagged with one copy of Atg11 with sfGFP and the other copy tagged with the HaloTag were treated with 40 nM JF646 HaloTag ligand for 30 min (*44*). The cells were then washed thrice for 15 min with CSM media to remove the unbound HaloTag ligand. The cells were then imaged for ∼1 h using a Leica DMi8 infinity TIRF inverted fluorescence microscope equipped with a Photometric Prime95B sCMOS camera, 100x 1.47 NA TIRF objective lens, 638 nm 150 mW laser module. Time-lapse movies of single molecules were acquired with a 15 ms time interval (total time points collected: 1000; exposure time: continuous acquisition with 15 ms camera processing time) and a 200 ms time interval (total time points collected: 200; exposure time: 50 ms).

For dwell time analysis, the particle tracking was performed (for 200 ms time interval movies) using the “TrackRecord” software developed in Matlab (The Matworks Inc.) (*44, 99, 100*). The software provides automated features for particle detection (using intensity thresholding), and tracking (using the nearest neighbour algorithm with molecules allowed to move a maximum of 6 pixels from 1 frame to the next, and only tracks that are at least 4 frames or longer are kept). Gaps in the tracks due to photoblinking of the fluorescent dye were closed upto 4 frames. The dwell time was determined by fitting the survival distribution (*99*) after photobleaching correction. Briefly, the survival histogram was generated from the time that each particle was stationary. In practice, even tightly bound particles move slightly due to chromatin and nuclear motion, and therefore a maximum frame-to-frame displacement of 470 nm (Rmin), and a two-frame displacement of 610 nm (Rmax) (both obtained from the motion of the chromatin-bound histone H3 (Hht1, (*101*)) have been used to define bound portions of each particle’s track. Because there is a chance that even a fast-diffusing molecule will move less than these thresholds, a further constraint on the minimum number of time points in the bound segment for each particle (Nmin) is used to reduce <1% the contribution of diffusing molecules to the survival histogram. The Nmin value used for 200 ms time-interval movies was 7. The total bound fraction is then calculated as the ratio of bound track segments to the total number of particles.

To extract residence times, the survival distribution, S(t), is fit by least squares to a mixed exponential decay with two rate constants, *k*_ns_=1/T_ns_ and *k*_s_ = 1/T_s_:

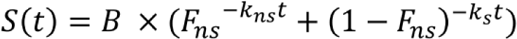

Where B is the bound fraction, and F_ns_ is the fraction of particles non-specifically bound. To check for over-fitting, the distribution is also fit to a single-component exponential:

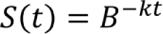

The fits are compared using an F-test to ensure that the two-component model gives a significant improvement over the single-component decay.

For quantifying diffusion parameters (fraction of bound/unbound molecules, diffusion coefficient), particle tracking was performed (from 15 ms time-interval movies) using DiaTrack Version 3.05 (Vallotton and Olivier 2013), with the following settings (*44*): remove blur 0.02, remove dim 50-250, maximum jump 6 pixels, where each pixel is 110 nm. This software determines the precise position of single molecules by Gaussian intensity fitting and assembles particle trajectories over multiple frames. The trajectory data exported from Diatrack was further converged into a single .csv file using a custom computational package ‘Sojourner’ (https://rdrr.io/github/sheng-liu/sojourner/). The Spot-On analysis was performed on three frames or longer trajectories using the web interface https://spoton.berkeley.edu/ (*45*). The bound fractions and diffusion coefficients were extracted from the Cumulative distribution function (CDF) of observed displacements over different time intervals. The cumulative displacement histograms were fitted with a 2-state model.

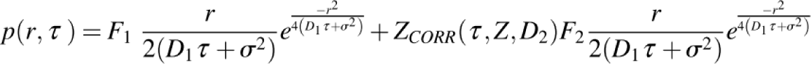

Where F_1_ and F_2_ are bound and free fractions, σ is single molecule localization error, D_1_ and D_2_ are diffusion coefficients of bound and free fractions, and Z_CORR_ is the correction factor for fast molecules moving out of axial detection range (*45*). The axial detection range for JF646 on our setup is 294 nm. The following settings were used on the Spot-On web interface: bin width 0.01, number of time points 6, jumps to consider 4, use entire trajectories – No, Max jump (µm) 1.5. For model fitting the following parameters were selected: D_bound_ (µm^2^/s) min 0.0001 max 0.5, D_free_ (µm^2^/s) min 0.5 max 5, F_bound_ min 0 max 1, Localization error (µm)-Fit from data-Yes min 0.01 max 0.1. Use Z Correction-No, Model Fit CDF, Iterations 3.

### Post-acquisition analysis

Spindle length was measured by tracking it using a straight line, and freehand (for bent spindle) tool in ImageJ after maximum intensity projection for each image. Spindle orientation was measured on images that were maximum intensity projected using the angle tool in ImageJ. The angle was calculated by measuring the smaller of the two angles that the spindle forms intersecting the mother-bud axis of the cell. The astral microtubules were measured after maximum intensity projection of the images using either a straight line or a freehand tool in ImageJ. The budding index was represented as the ratio of the diameter of the daughter cell to the diameter of the mother cell. Throughout the manuscript, large-budded cells with a budding index of >0.6 are taken into consideration for all the analyses (*86*). Statistical analyses were done using GraphPad Prism 8.4.0 software. Student’s *t*-test followed by Unpaired *t*-test for comparing two groups. The data sets represented as bar graphs were analysed using either One-way ANOVA or two-way ANOVA for multiple groups and have been mentioned in the figure legends where applicable. Scatter plots with data from three independent biological repeats were plotted as SuperPlots (*102*), displaying individual colour-coded data. The statistical analyses were done with three independent biological replicates.

## Acknowledgements

We thank J. Heitman, L. Sreekumar, S. Sridhar, V. Yadav, and K. Guin for the critical reading of the manuscript. We thank D. J. Klionsky, C. Kraft, S. K. Ghosh, S. Bhattacharyya, and F. Monje-Casas for sharing yeast strains and vectors. We acknowledge A. Desai (UCSD, California), G. Dey (EMBL, Heidelberg), G. Pereira (University of Heidelberg), E. Schiebel (DKFZ-ZMBH, Heidelberg), S. Laxman (InStem, Bengaluru) and U. Surana (A*STAR, Singapore) for the valuable suggestions. We thank M. Sirajuddin (InStem, Bengaluru) for the valuable discussion related to aMT dynamics. We also acknowledge B. Bhojappa (IISc, Bengaluru) for providing Cdc14-GFP Tub1-mCherry tagged strain and helping with the imaging and S. Palani (IISc, Bengaluru) for the constructive suggestions. We thank B. Suma and K. Jayaram at the imaging facility at JNCASR. Financial support from the DBT-RA Program in Biotechnology and Life Sciences is gratefully acknowledged by MHR. MHR also acknowledges EMBL Corporate Partnership Programme Fellowship and EMBO Scientific Exchange Grant 10212. The award of the JC Bose Fellowship (JCB/2020/000021), SERB and JNCASR intramural funding support to KS is acknowledged. This work was also supported by SERB grant (CRG/2019/004892) to RM. JV was supported by intramural financial support from JNCASR. RA is supported by intramural financial support from JNCASR. GM lab is supported by the Har-Govind Khorana Innovative Young Biotechnologist Award (BT/13/IYBA/2020/10), Ramalingaswami fellowship (BT/RLF/Re-entry/53/2020) from the Department of Biotechnology, Govt. of India and JICA (Japan International Cooperation Agency). NKP acknowledges the Prime Minister’s Research Fellowship (PMRF ID: 2001700) for the financial support. We also thank the members of the Molecular Mycology Laboratory and autophagy lab for their valuable inputs and comments. Some images were created with BioRender.com.

## Author Contributions

KS conceived the idea; MHR and KS designed experiments; MHR performed the majority of the experiments; RA created strains used in the BiFC assays, yeast-two hybrid assays, and western blot to study Pds1 levels; JV performed the initial screen of autophagy mutants involved in chromosome segregation, stability of monocentric plasmid and *CEN* transcriptional read-through assay; NKP and GM performed single-molecule tracking experiments and analyzed data; RC carried out *in silico* protein structure-function analysis; MHR, RA, JV, RC, and RM contributed new reagents/analytical tools; MHR, RA, JV, RM, GM, and KS analyzed the data; MHR, RA, GM, and KS wrote the manuscript and all authors contributed in editing the manuscript; KS gathered the majority of the funding and supervised the work.

## Supporting Tables

**Supporting Table S1:**
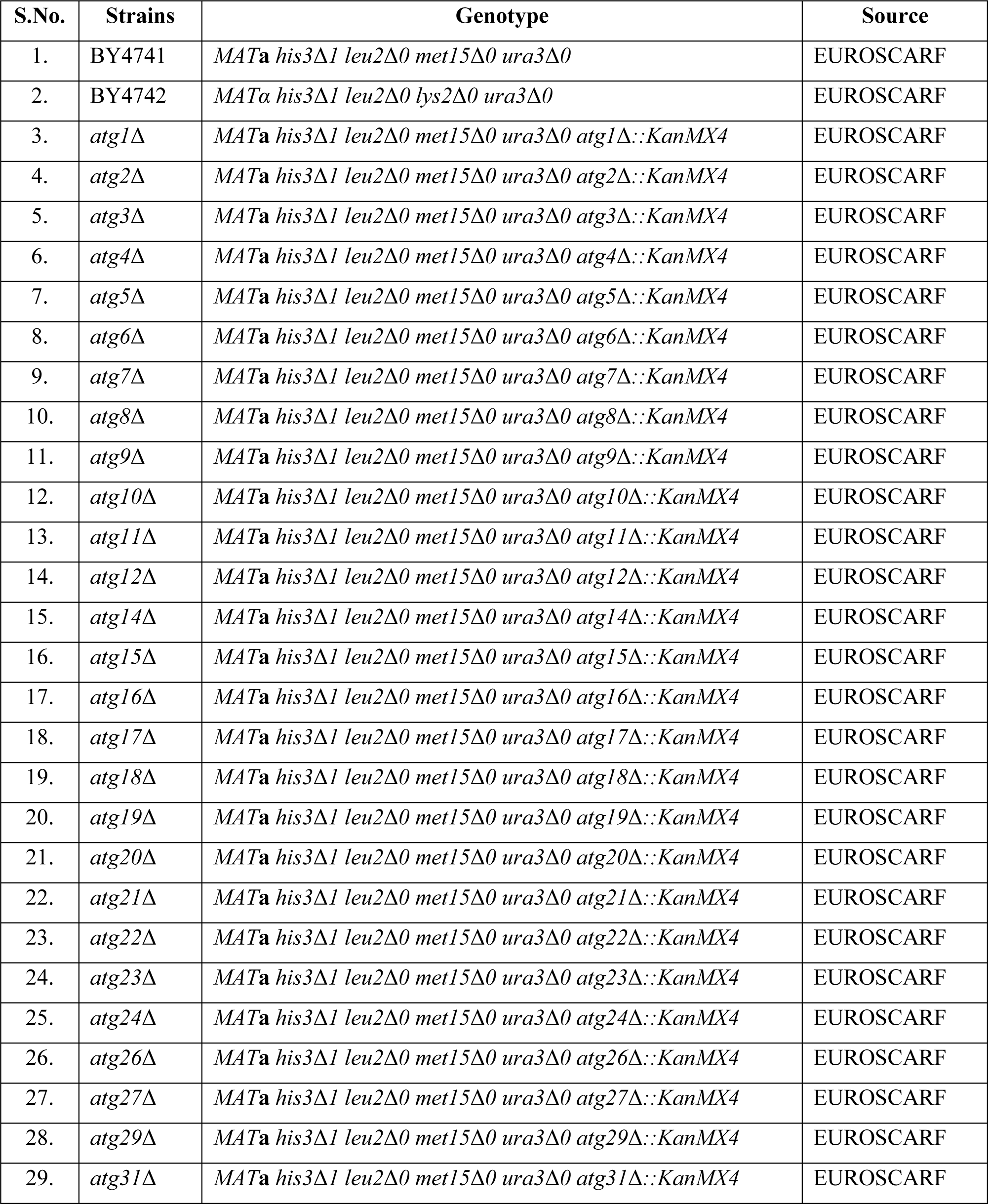
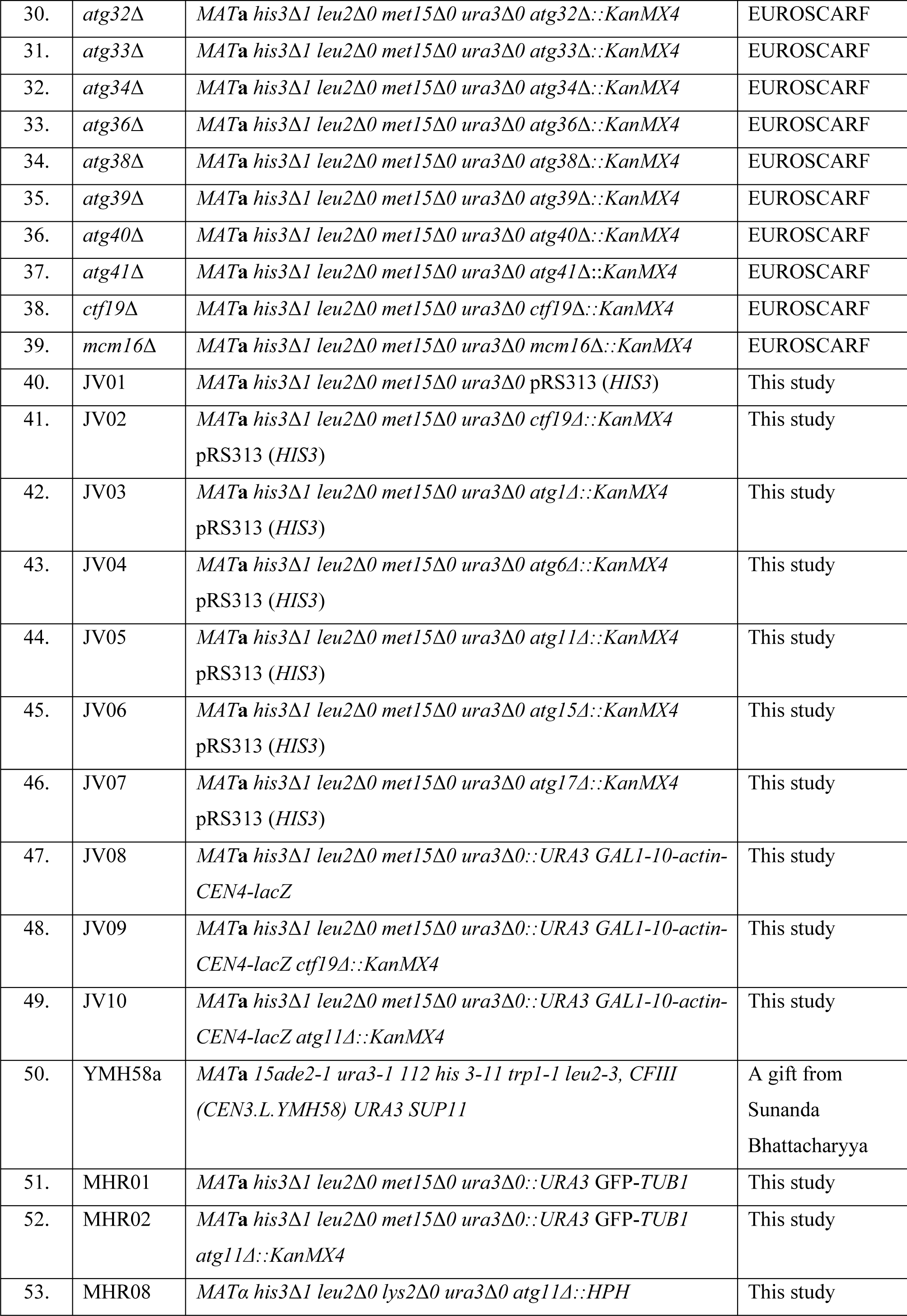
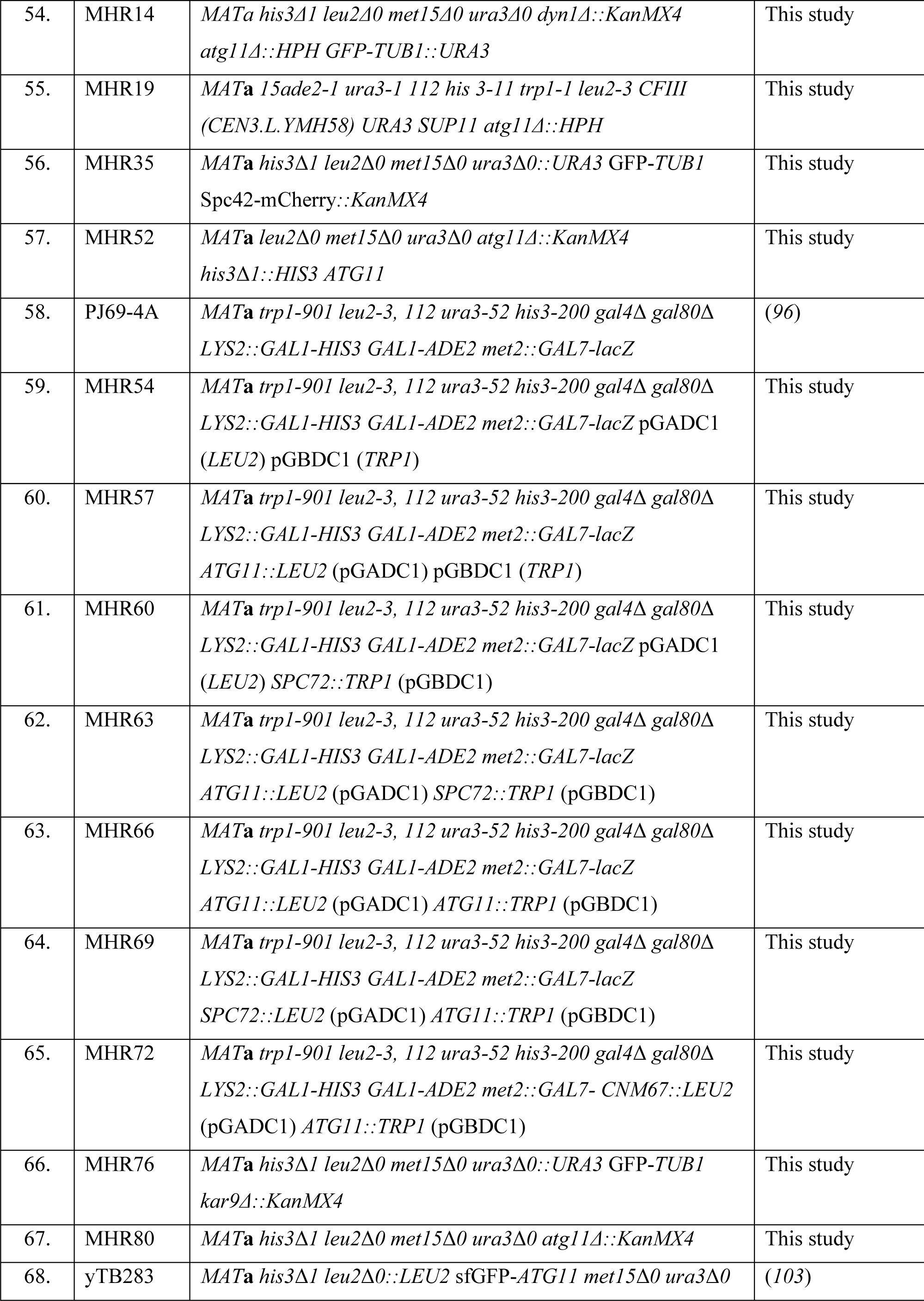
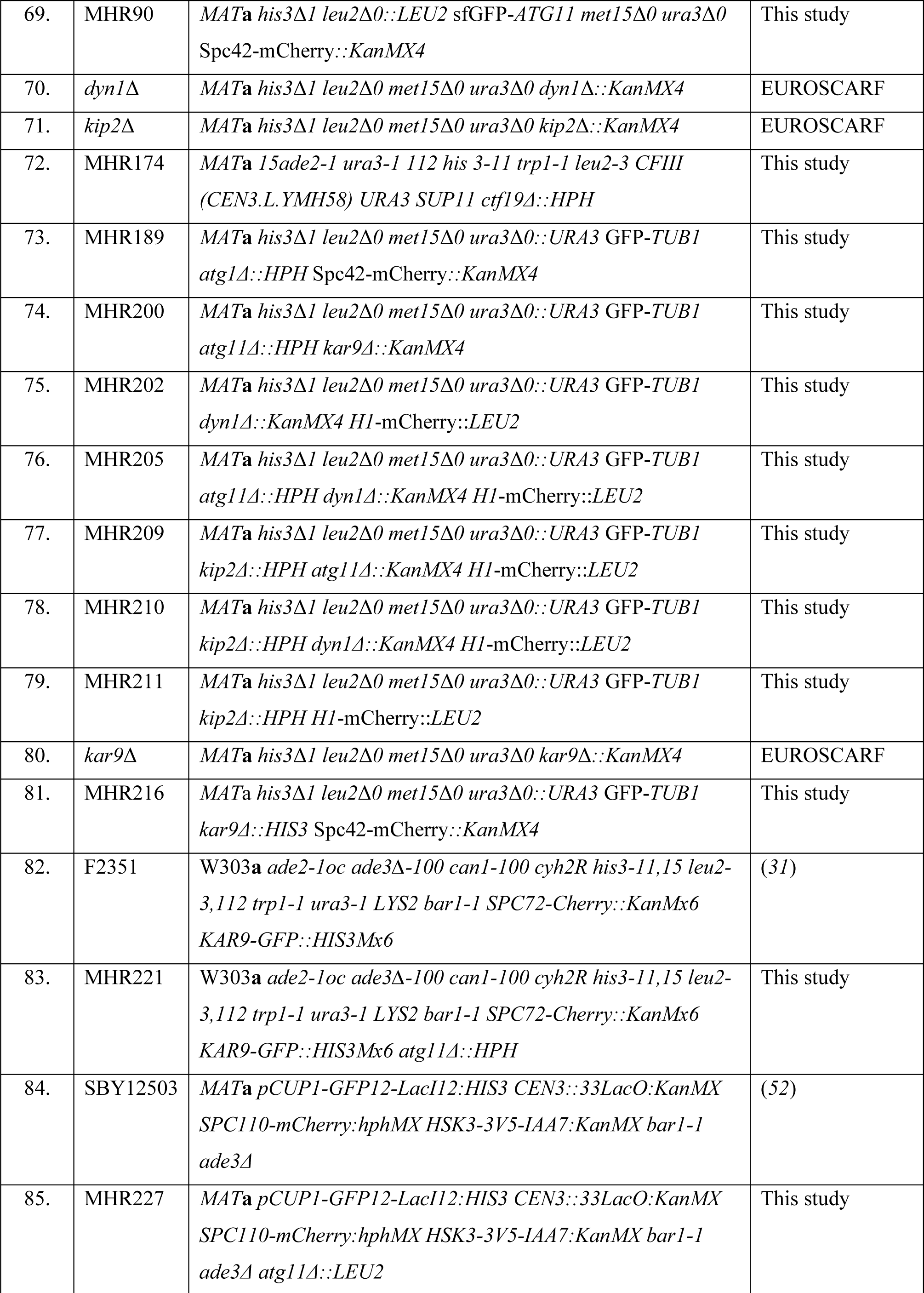
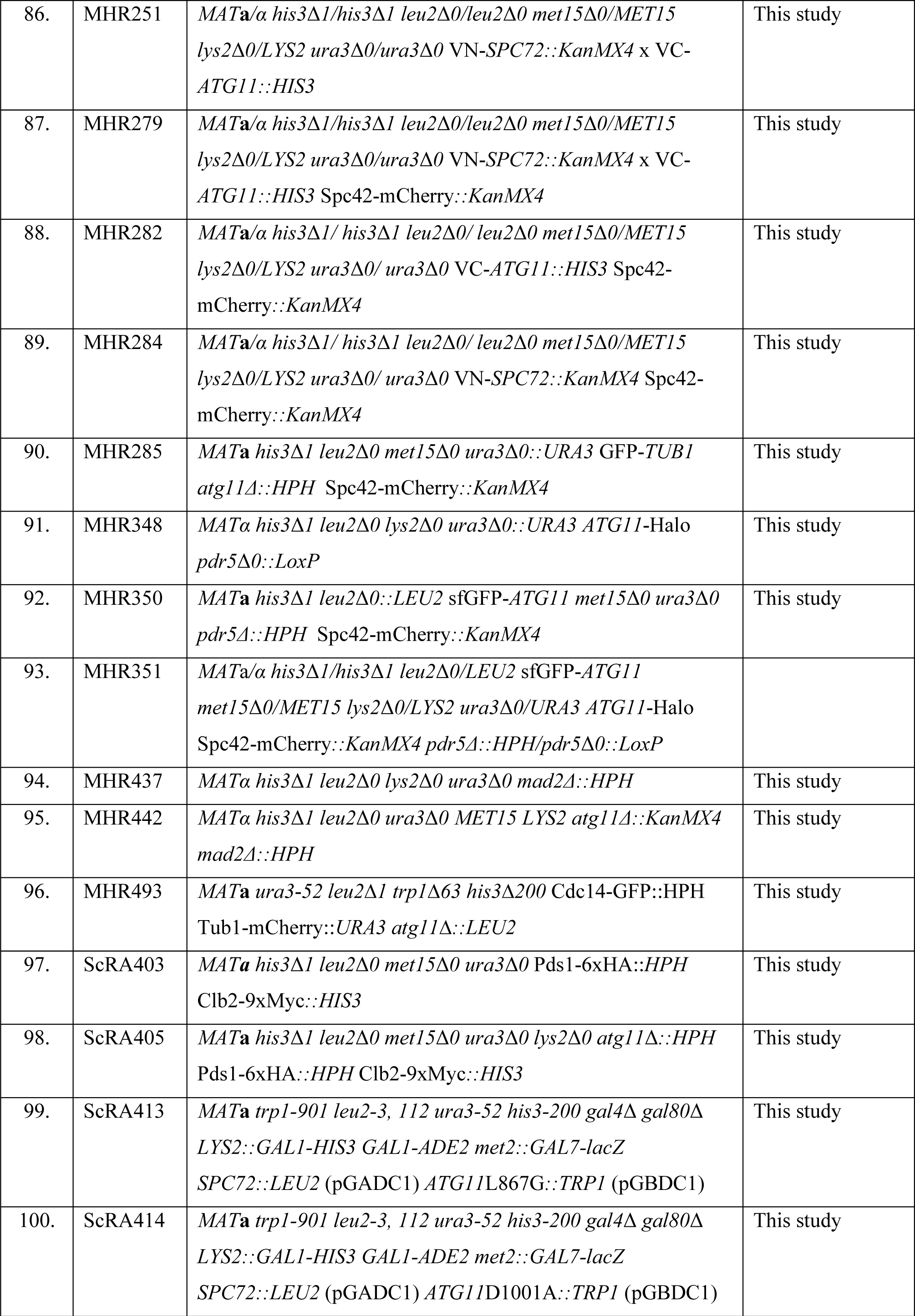
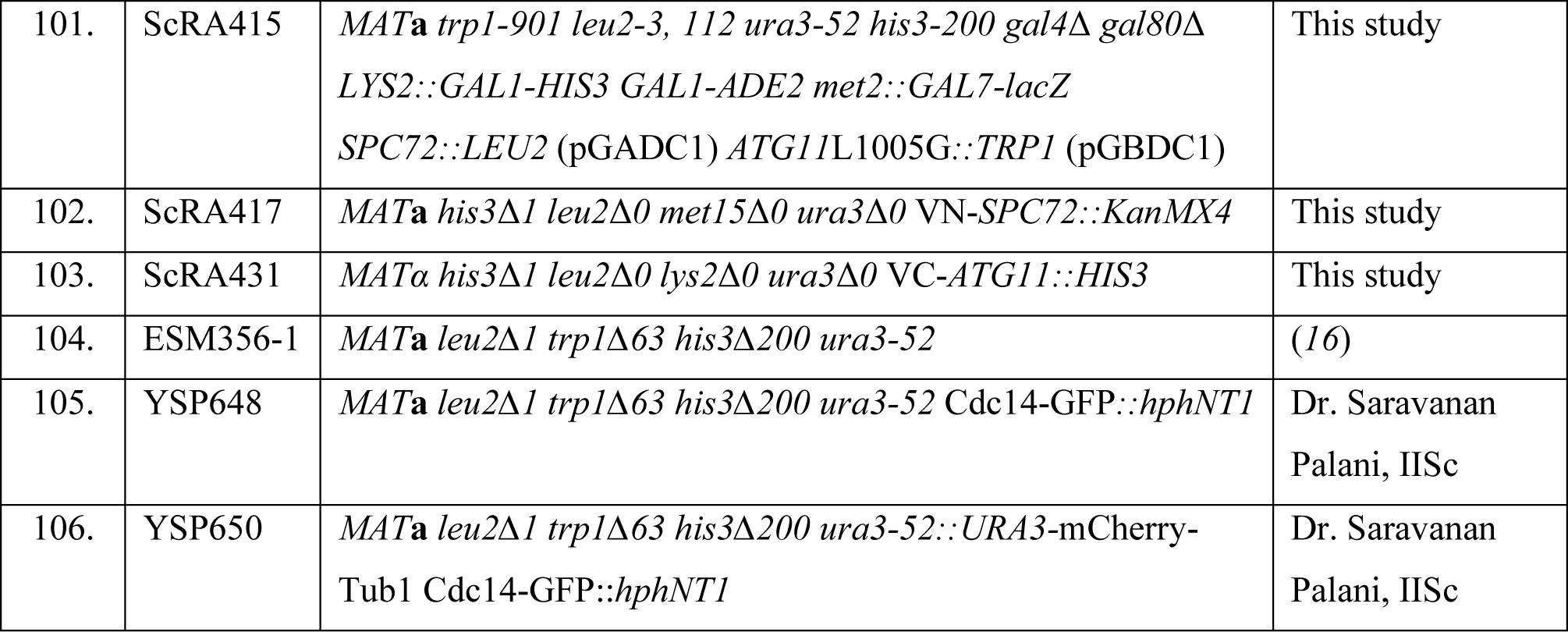
*S. cerevisiae* strains used in this study.

**Supporting Table S2:**
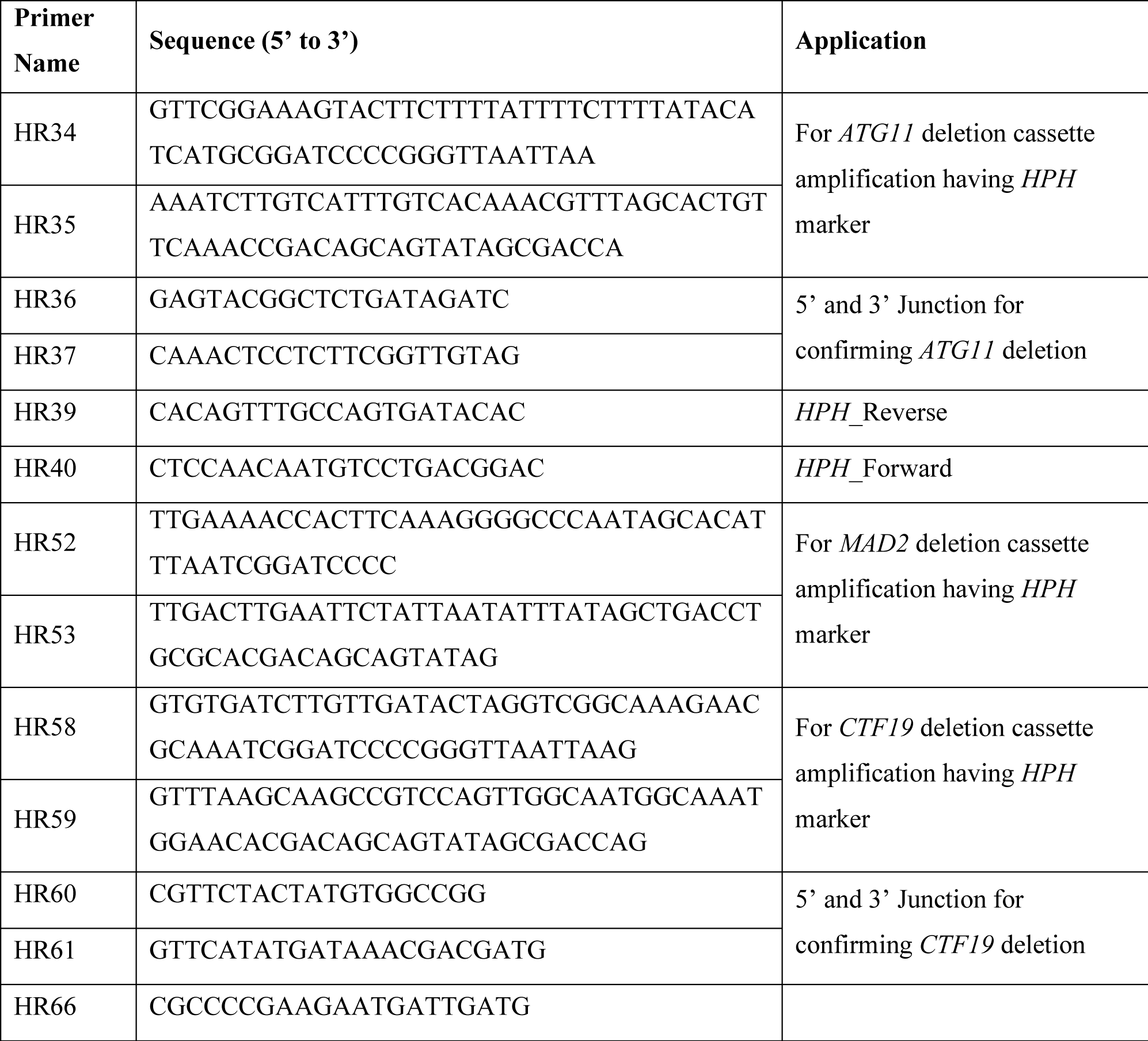
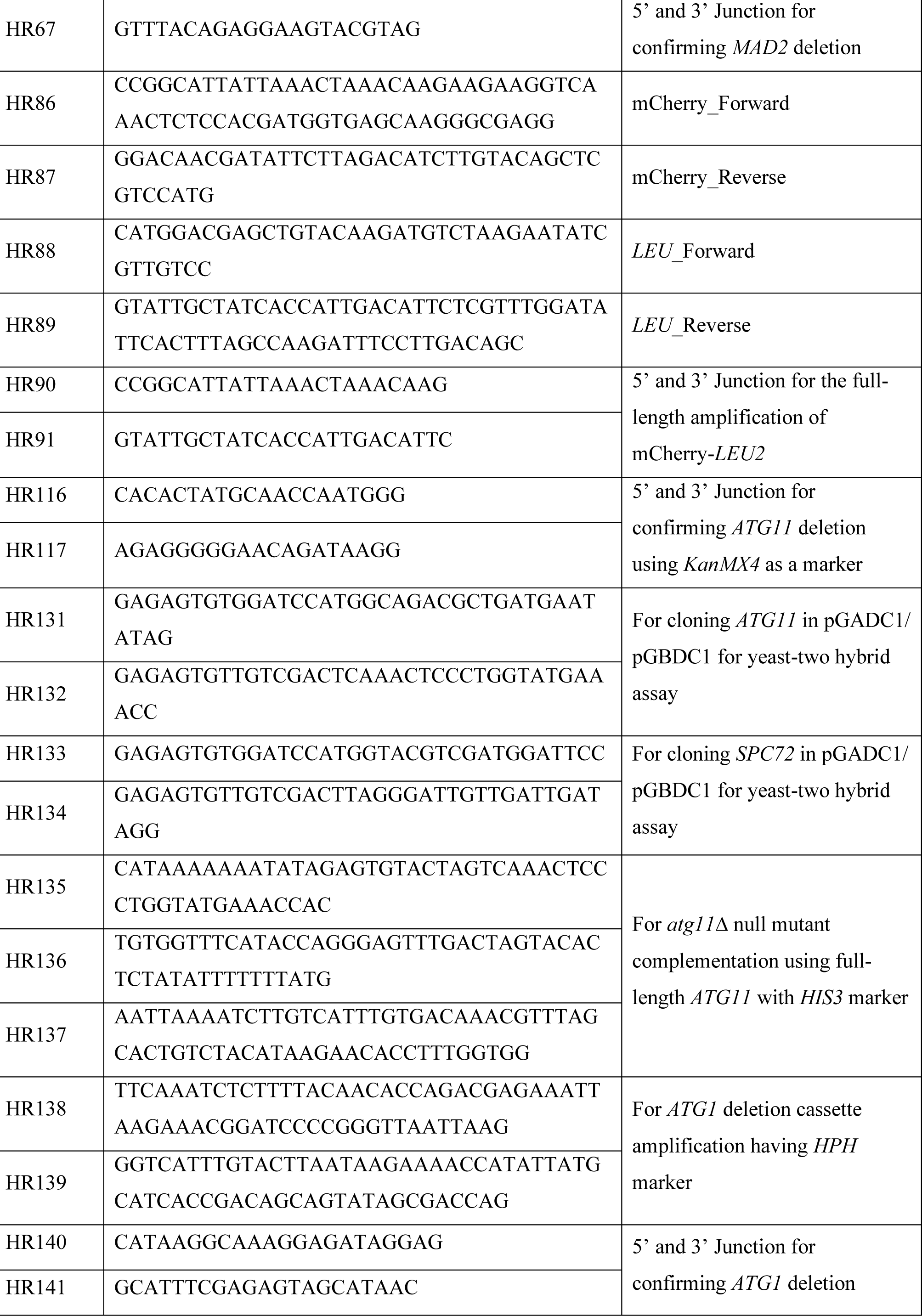
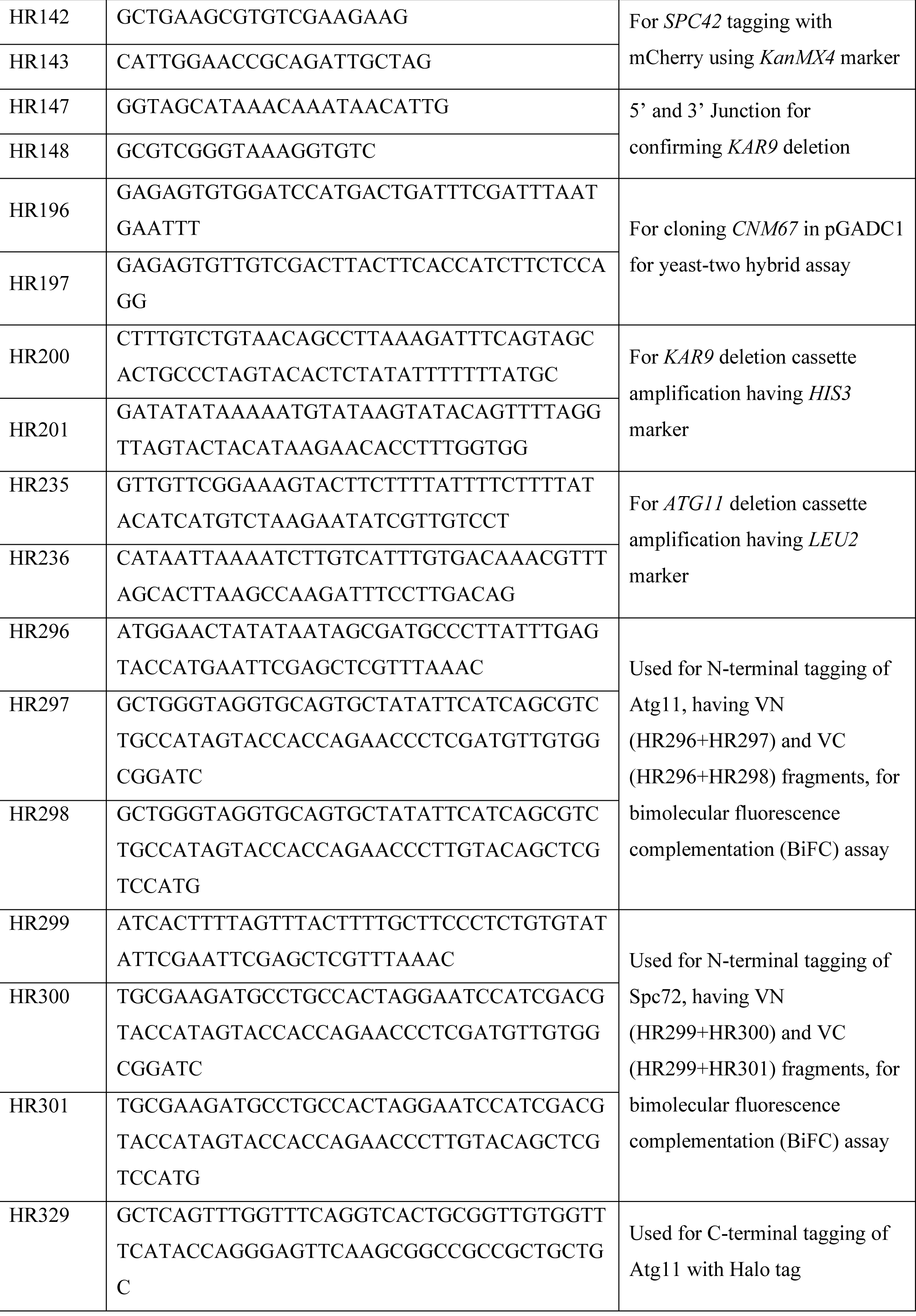
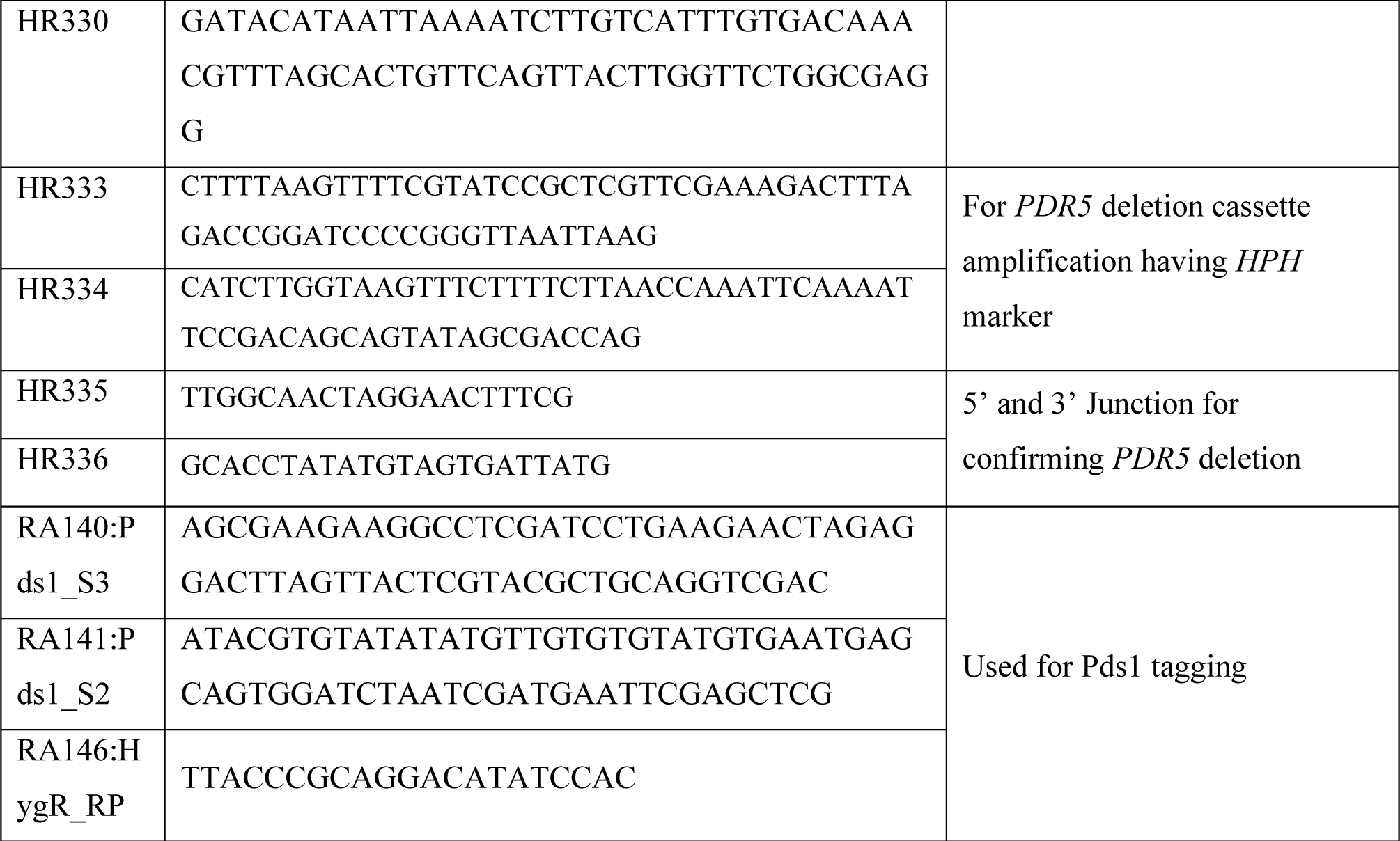
Oligonucleotides used in this study.

**Supporting Table S3:**
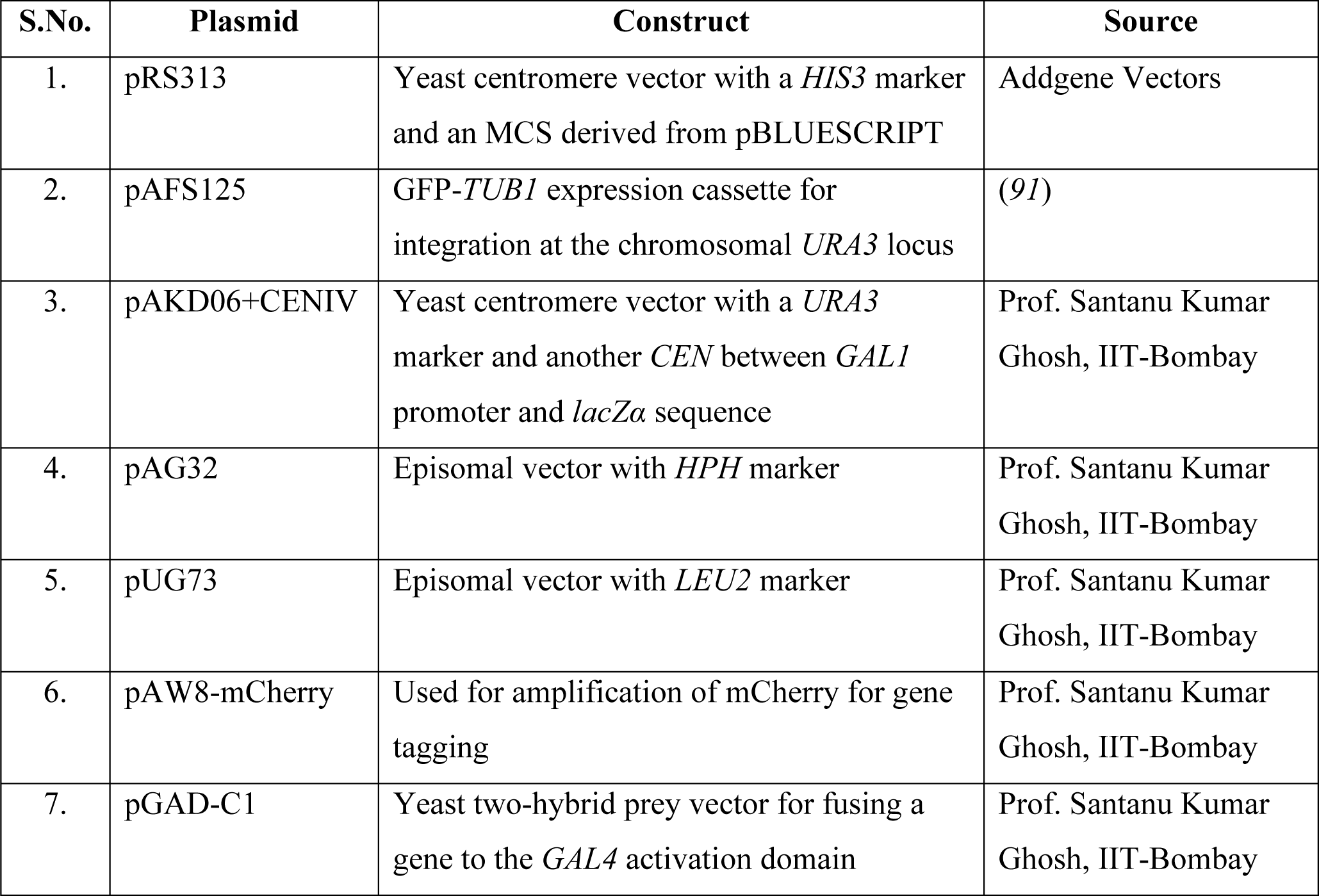
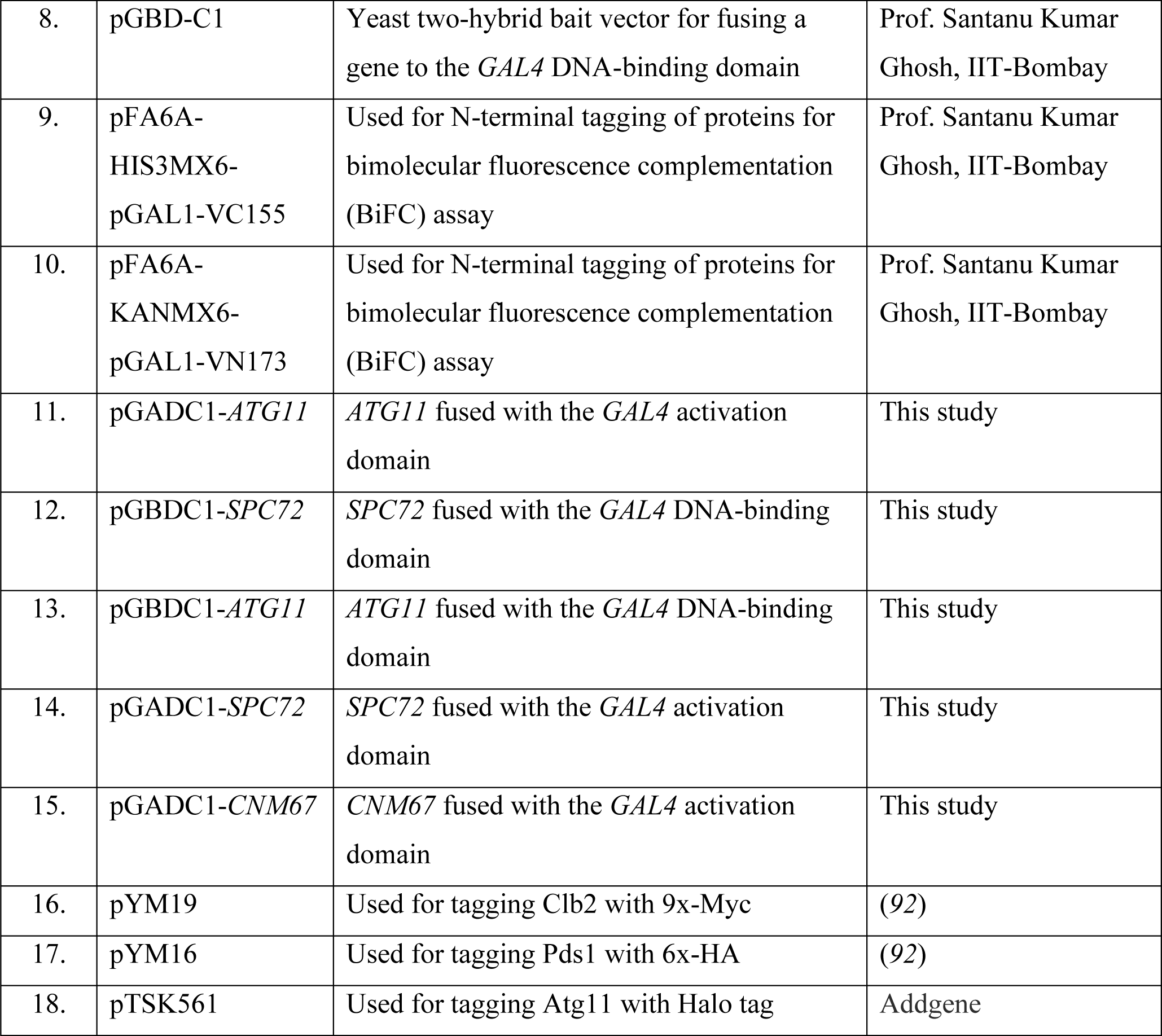
Plasmids used in this study.

